# Astrocyte-induced firing in primary afferent axons

**DOI:** 10.1101/2024.06.07.597942

**Authors:** Fanny Gaudel, Julia Giraud, Philippe Morquette, Marc Couillard-Larocque, Dorly Verdier, Arlette Kolta

**Author notes:** **Corresponding Author:** (AK). F. M. Kirby Neurobiology Center and Department of Neurology, Boston Children’s Hospital and Harvard Medical School. Boston, MA 02115, USA. These authors contributed equally to this work.

## Abstract

The mesencephalic trigeminal nucleus is unique in that it contains the cell bodies of large-caliber primary afferents that are usually located in the periphery in the dorsal root ganglia or trigeminal ganglia. The activity of these afferents is typically associated with proprioception of the jaw-closing muscles or mechanoreception on the teeth and periodontal ligament. However, like other large-caliber afferents from the body which display ectopic firing in neuropathic pain models, these afferents exhibit increased excitability and ectopic discharges even in a relatively mild muscle pain model. These discharges normally emerge from subthreshold membrane oscillations (SMOs) supported by a persistent sodium current (*I*_NaP_) which is exquisitely sensitive to extracellular Ca^2+^-decreases. We have shown in the trigeminal main sensory nucleus that the release of a Ca^2+^-binding astrocytic protein, S100β, is sufficient to modulate this sodium current. Here, we explore if this astrocyte-dependent mechanism contributes to emergence of this hyperexcitability and aim to localize the cellular site where ectopic discharge may arise using whole-cell patch-clamp recordings, confocal imaging, and immunohistochemistry methods on mice brain slices. We found that astrocytes, by lowering [Ca^2+^]_e_ at focal points along the axons of NVmes neurons through S100β, enhance the amplitude of the Na_V_1.6-dependent SMOs leading to ectopic firing. These findings suggest a crucial role for astrocytes in excitability regulation and raise questions about this neuron-astrocyte interaction as a key contributor to hyperexcitability in several pathologies.

## Introduction

The primary afferents (PAs) innervating the spindles of the jaw-closing muscles and the pressoreceptors of the periodontal ligaments have their somata in the mesencephalic trigeminal nucleus (NVmes), located centrally in the brainstem, where they receive inputs from several structures in the forebrain, mid-brain (1–3), and lower brain stem (4–7). These neurons exhibit voltage-dependent rapid subthreshold membrane oscillations (SMOs) which greatly influence cellular excitability since they often lead to repetitive firing. Both, the SMOs and repetitive firing properties of NVmes neurons, rely on a persistent sodium current (*I*_NaP_) (8–12) and evidence of a strong contribution of Na_V_1.6 channels to *I*_NaP_ and SMOs has been shown in these neurons (8). However, it is known that the Na_V_1.6 channels distribution is not uniform on the membrane of myelinated neurons: a higher concentration is found at the axon initial segment and the nodes of Ranvier (13–18) where they contribute to the action potential initiation and propagation, respectively. This implies that in NVmes neurons, both activities may arise from their axon membrane electrical properties rather than the soma. Therefore, local modulation of axonal Na_V_1.6 could have profound effect on these neurons output.

Variation of extracellular calcium concentration ([Ca^2+^]_e_) modulates voltage-gated sodium channels (19) and *I*_NaP_ in particular is enhanced by the decrease of [Ca^2+^]_e_. Our team has shown that astrocytes can play a key role in this mechanism through release of the Ca^2+^-binding protein, S100β, which by decreasing [Ca^2+^]_e_ and enhancing *I*_NaP_, changes the discharge pattern of neurons in the trigeminal main sensory nucleus (NVsnpr) (20) and integration properties of layer 5 pyramidal neurons in the visual cortex (21). We postulate that this neuron-astrocyte interaction occurs at a precise neuronal compartment in the NVmes where Na_V_1.6 channels are enriched. We hypothesize that it could explain the reported excitability changes of the large-diameter primary afferents in various pain models. For example, in the acid-induced jaw muscle chronic myalgia model (22) in which increased amplitude of the SMOs and spontaneous ectopic firing were observed, despite a hyperpolarizing shift in their firing and SMOs threshold. Thus, using a combination of whole-cell patch clamp recordings, local applications of Ca^2+^-chelating agents, immunohistochemistry, and optogenetic activation of peri-axonal astrocytes, we tested whether manipulation of astrocytes or local decreases of [Ca^2+^]_e_ along the axons of NVmes neurons affect the SMOs and repetitive firing capability of these neurons through an S100β/Na_V_1.6-dependent mechanism. We found that locally chelating the calcium around the axons of NVmes neurons increases the SMOs amplitude and hyperpolarizes the SMOs and firing thresholds thereby increasing the occurrence of repetitive firings in these neurons. We also found that astrocytes are closely positioned next to the Na_V_1.6 immunopositive axons of the NVmes neurons and that their optogenetic activation enhances Na_V_1.6-dependent SMOs leading to ectopic firing through the release of S100β along the axons of NVmes neurons.

## Results

### Electrophysiological properties of NVmes neurons

160 neurons recorded in the mesencephalic trigeminal nucleus (NVmes) of 44 WT mice and 68 GFAP-ChR2 mice fulfilled the inclusion criteria. The recorded neurons were filled with Alexa Fluor (488 or 594), and all showed the typical pseudo-unipolar morphology of primary sensory afferents consisting of a spherical or ovoid cell body attached to a single process. Their basic electrophysiological characteristics are summarised in **Table 1**. There were no statistical differences in the resting membrane potential, the input resistance, and the firing threshold between neurons recorded from WT and GFAP-ChR2 mice (**Table 1**, Kruskall-Wallis test, P ˃ 0.05). Therefore, all the data obtained from both mice lines were pooled together. The pooled neurons had a resting membrane potential (RMP) of -54 ± 0.2 mV, a firing threshold of -43 ± 0.4 mV, and an input resistance of 82 ± 4 MΩ. All the recorded neurons showed a strong inward rectification producing a prominent sag (**Fig 1A**, arrow) upon membrane hyperpolarization, typical of these cells, while 96 (60%) of them fired a rebound action potential at the offset of the hyperpolarizing pulses (**Fig 1A**, arrowhead). Accommodation of firing upon membrane depolarization occurred in 120 of the 160 (75%) neurons, with 68 (57%) of them discharging only a single action potential (**Fig 1A**), even with long-duration (up to 1000 ms) pulses. The I-V curve shown in **Figure 1A** is typical of the accommodating NVmes neurons and reveals an outward rectification during depolarization and an inward rectification during hyperpolarization, both emphasized by the linear fitting (red straight line) of the non-rectifying portion of the I-V curve. The 40 (25%) cells that did not show firing accommodation either fired recurrent bursts (n=30) as illustrated in **Figure 1B** or fired without pause for the duration of the depolarizing pulses (n=10; **Fig 1C**). Thus, the recorded NVmes neurons in mice could be classified into the same three subtypes reported by Yang *et al*. (23) in rats: the spike-adaptative, the burst-firing, and the tonic-firing neurons. All the burst-firing neurons exhibited voltage-dependent subthreshold membrane potential oscillations (SMOs, inset in **Fig 1B**, right) while only 32 (27%) out of the 120 adaptative neurons exhibited such activity. SMOs, most of the time, could not be sampled in the tonic-firing neurons since they tend to fire repetitively when depolarized. The mean frequency of this oscillatory activity in 30 burst-firing and 32 spike-adaptative NVmes neurons was 82 ± 3 Hz, their averaged peak-to-peak amplitude was 1.5 ± 0.1 mV and they appeared at an average threshold membrane potential of -45 ± 0.6 mV as determined by a ramp current injection protocol. As already shown in numerous studies in rats (10, 24) and mice (25), these SMOs were suppressed by membrane hyperpolarization (**Fig 1D**, bottom trace), and increased in amplitude with membrane depolarization until strong repetitive firing was elicited (**Fig 1D**, top trace). When measured in bursting neurons in which clear SMOs could be seen next to a burst (n=25), the SMOs frequency in each cell (**black** dots in graph in **Fig 1D**) paralleled the intra-burst firing frequency (**grey** dots in graph in **Fig 1D**) and were significantly correlated (linear regression, r = 0.96, P ˂ 0.001), indicating that these SMOs are a key factor that contributes to greater cellular excitability. The SMOs are supported by *I*_NaP_ (10, 24, 25), and we as others (23), have hypothesized that any factor that impacts *I*_NaP_, will indirectly modulate the excitability of these neurons.

**Fig 1.**
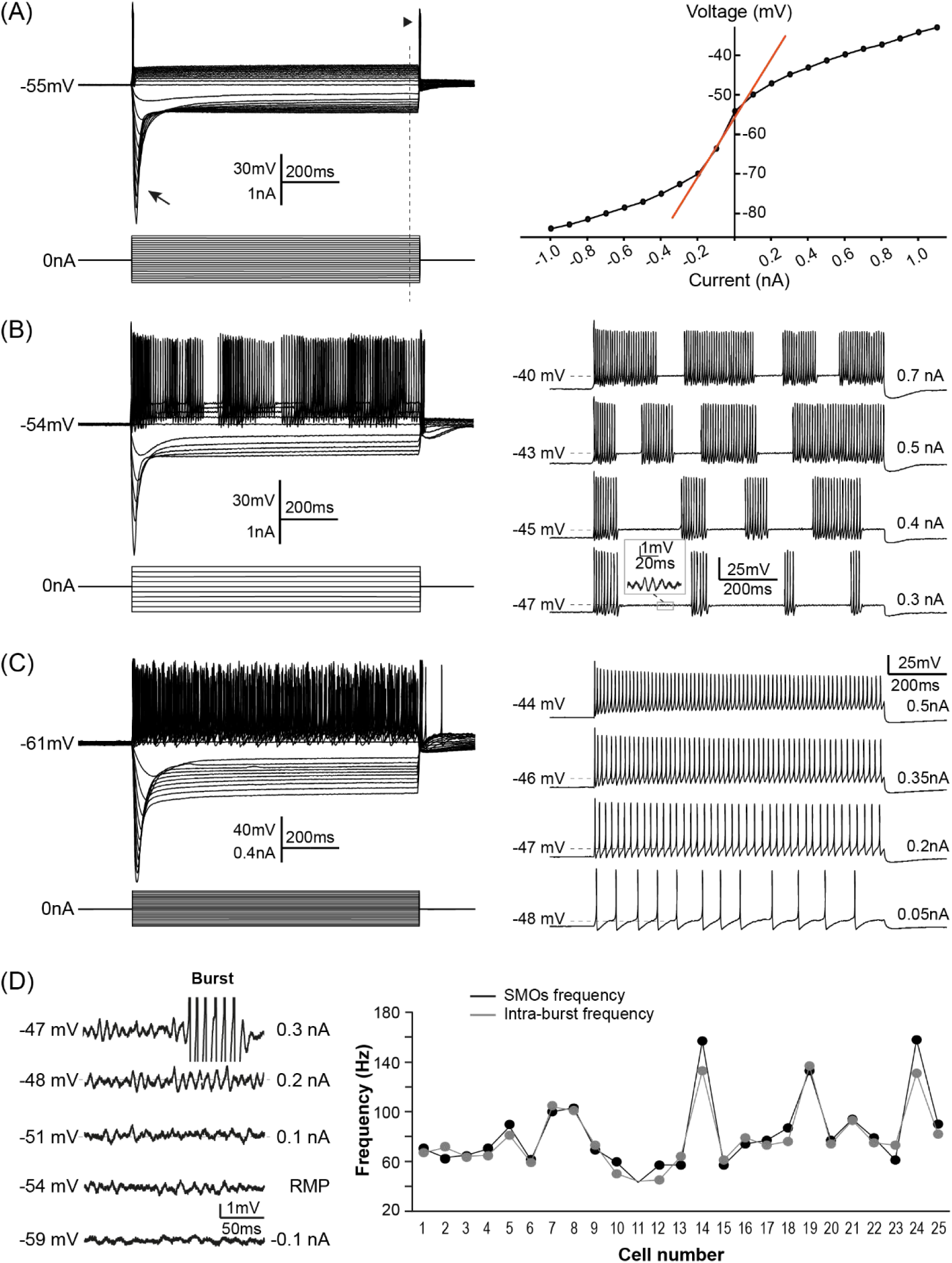
Diversity of NVmes neurons firing profiles. Left, top traces: Membrane responses of a spike-adaptative **(A)**, a burst-firing **(B)**, and a tonic-firing **(C)** NVmes neurons to injections of hyperpolarizing and depolarizing current pulses (bottom traces). The arrow and arrowhead point the sag and the rebound action potential, respectively. The vertical dotted line indicates the position of the membrane voltage and the current injection lecture for the I-V curve. **(A) Right**: I-V curve showing the inward and outward membrane rectification of a spike-adaptative neuron in response to hyperpolarizing and depolarizing injected current, respectively. The red line is the linear fitting to the non-rectifying portion of the I-V curve. (**B, C) Right**: isolated traces to distinguish bursting and tonic firing profiles. Inset in **(B, right)** shows SMOs in a burst-firing NVmes neuron. **(D) Left**: Membrane voltage recordings showing the effect of membrane potential on the SMOs. They are abolished with membrane hyperpolarization (**bottom** trace), increased, and lead to firing with membrane depolarization (**top** trace). **(D) Right**: Plot of the intra-burst firing frequency (**grey** dots) and SMOs frequency (**black** dots) in 25 NVmes neurons at the bursting threshold for each cell showing how they parallel each other.

**TABLE 1.** Electrophysiological characteristics of NVmes neurons.

| Electrophysiological characteristics | WT mice<br>N = 61 | GFAP-ChR2 mice<br>N = 99 | WT + GFAP-ChR2 mice<br>N = 160 | Nav1.6 KO mice<br>N = 15 |
| --- | --- | --- | --- | --- |
| RMP (mV) | -55 ± 0.5 | -54 ± 0.3 | -54 ± 0.2 | -56 ± 0.9 |
| Firing threshold (mV) | -42 ± 0.7 | -43 ± 0.4 | -43 ± 0.4 | -37 ± 1.8 |
| Input resistance (MΩ) | 83 ± 8 | 82 ± 5 | 82 ± 4 | 70 ± 8 |

Abbreviations: ChR2, Channelrhodopsin 2; KO, Knock-out; NVmes, mesencephalic trigeminal nucleus; N, number of neurons; RMP, resting membrane potential; WT, wild type.
Values are mean ± SEM.
\* $P = 0.03$ , Kruskal Wallis test with Bonferroni correction.
$P > 0.05$ for comparisons of the 3 parameters between WT and GFAP-ChR2 mice, Kruskal Wallis test.

### Axonal applications of BAPTA cause firing in NVmes neurons

Our previous work, using whole-cell recordings of NVsnpr and cortical neurons, has demonstrated the ability to modulate *I*_NaP_ or *I*_NaP_-dependant neuronal activity by producing local decreases of Ca^2+^ with local applications of Ca^2+^-chelators such as BAPTA or S100β (20, 21). Using a similar approach here to test the effect of *I*_NaP_ modulation on the excitability of NVmes neurons, BAPTA was locally applied near the soma or the stem axon of 40 NVmes neurons from 29 mice. To alleviate the text, all the numbers related to the effects of these applications are reported in **Table 2**. The only observed effect of BAPTA applications near the soma was a long-lasting hyperpolarization that outlasted the duration of the BAPTA applications by about 13 s (**Table 2**, **Fig 2A**, left trace). Similar hyperpolarization was also observed following axonal BAPTA applications but only in 13% of cases (**Table 2**, **Fig 2B**, top left trace). Depolarization was observed only once (not shown), and no effect were seen with 6 of the 69 applications. Strikingly, the most prevalent effect (77% of cases) of locally applied BAPTA near the axon was induction of firing (**Table 2**, **Fig 2B**, left traces). In most cases (n= 36/53) the firing overrode a depolarizing plateau of 1.9 ± 0.1 mV that occurred 0.4 ± 0.1 s after the onset of the application (**Fig 2B**, second left trace). In seven cases, the firing was either preceded by or occurred concomitantly with a membrane potential hyperpolarization of 1.1 ± 0.7 mV starting 0.2 ± 0.04 s after the onset of the application (**Fig 2B**, third left trace). In the remaining 10 cases, the firing occurred directly without prior membrane potential change at a latency of 1.6 ± 0.9 s (**Fig 2B**, bottom left trace). Four different firing patterns were elicited by these axonal BAPTA applications: high-frequency trains (60 ± 7 Hz, n=16, **Fig 3A**, top trace); recurrent bursting (inter-burst frequency: 5.5 ± 0.6 Hz, intra-burst frequency: 72 ± 6 Hz, n=11, **Fig 3A**, second trace); low-frequency trains (12 ± 4 Hz, n=7, **Fig 3A**, third trace), and a mix of bursting and high- or low-frequency trains (n= 12 and 7 respectively, **Fig 3A**, fourth and fifth traces respectively). The vertical bar charts in **Figure 3B** and **3C** illustrate the relative distribution of the different responses and the different firing patterns, respectively, observed in response to BAPTA (blue bars) application near the soma or axonal process of the recorded neurons.

**Fig 2.**
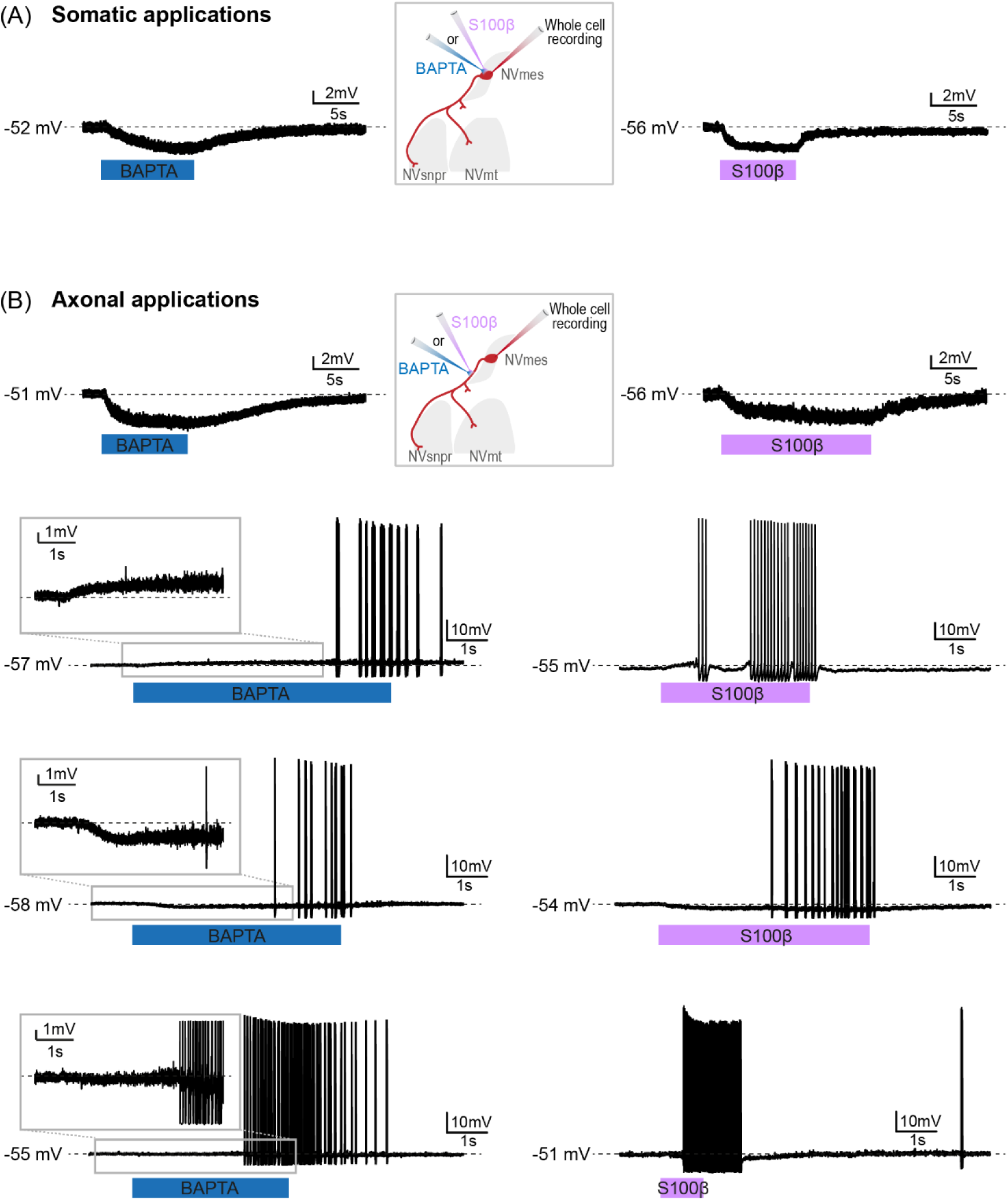
Chelation of extracellular calcium exerts a variety of effects on NVmes neurons. **(A):** Somatic applications of Ca^2+^ chelators BAPTA (**left** trace) and S100β (**right** trace) induce a slight hyperpolarization of NVmes neurons’ membrane potential. **(B):** Applications of Ca^2+^ chelators BAPTA (**left** traces) or S100β (**right** traces) along the axonal process of NVmes neurons induce cell hyperpolarization (**top left and right** traces, respectively) or firing, which could be preceded by membrane depolarization (**second left and right** traces), hyperpolarization (**third left and right** traces), or no change in membrane potential (**bottom left and right** traces). **Middle Insets**: Cartoons illustrating the experimental set-ups. **Left Insets**: Zooms on the initial part of the firing responses. All somatic applications resulted in cell hyperpolarization, while most axonal applications triggered cell firing. *Abbreviations: NVmes, mesencephalic trigeminal nucleus; NVmt, trigeminal motor nucleus; NVsnpr, trigeminal main sensory nucleus*.

**Fig 3.**
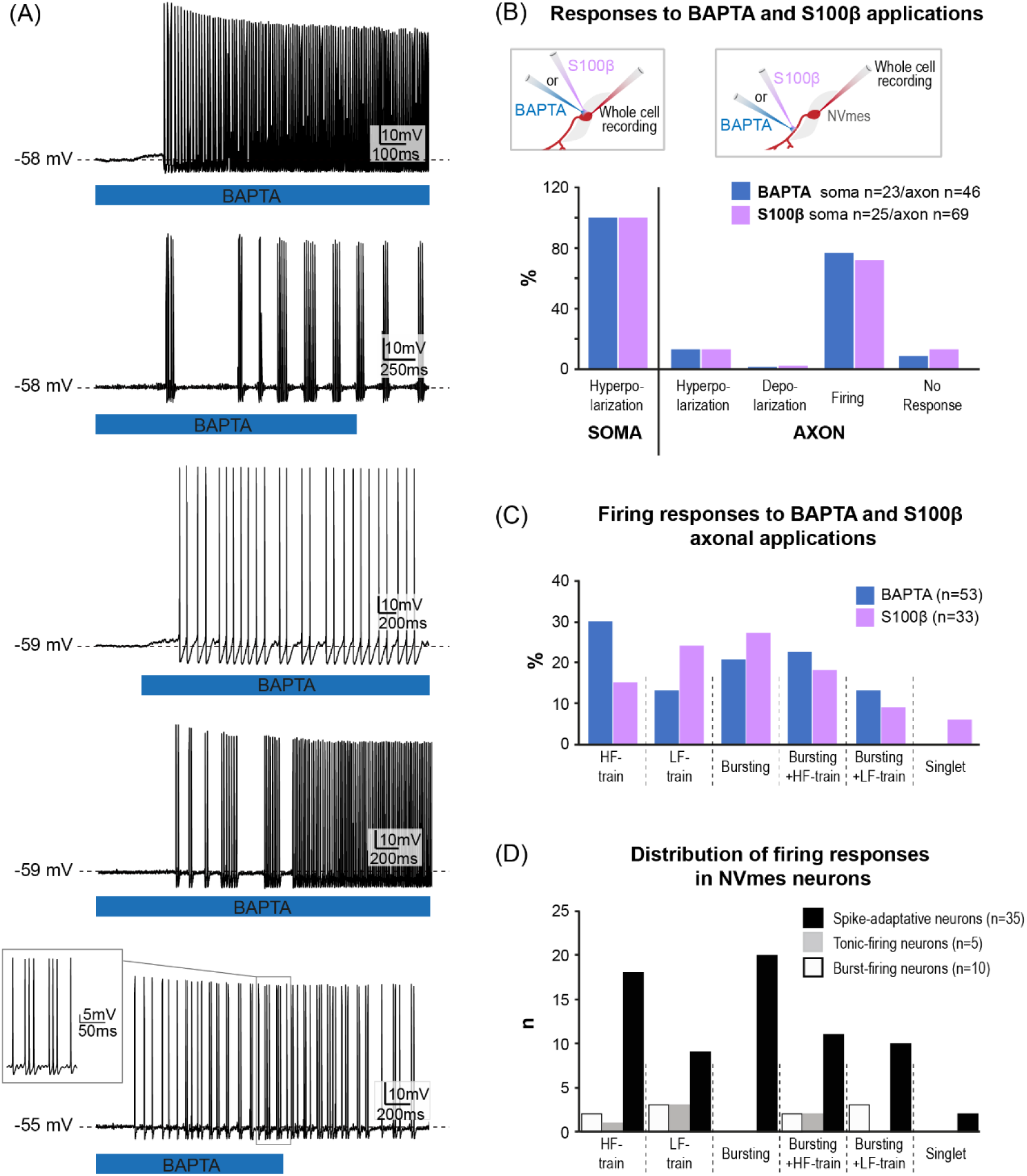
Patterns of axonal BAPTA- or S100β -induced firing in NVmes neurons. The axonal applications of the Ca^2+^ chelator BAPTA induced a train of high-frequency repetitive firing (**A, top left** trace) in a majority of NVmes neurons. In some neurons, BAPTA triggered a mix of neuronal bursting and train of action potentials (**A, top righ**t trace), bursting only (**A, middle left** trace), bursting associated with single action potentials (**A, middle right** trace), or a train of low-frequency single action potentials for the duration of BAPTA application (**A, bottom left** trace). **(B):** Bar chart quantifying the response profiles of NVmes neurons to somatic or axonal applications of BAPTA (**blue**), or of S100β (**mauve**) in WT/GFAP-ChR2 mice. **(C):** Bar chart of the relative distribution of neuronal firing types elicited after axonal applications of BAPTA (**blue** bars), as illustrated in **(A)**, and S100β (**mauve** bars, traces not shown in A). **(D):** Bar chart of the relative distribution of the neuronal firing responses elicited after axonal applications of BAPTA and S100β in the spike-adaptative (black bars), tonic-firing (grey bars), and burst-firing (empty bars) NVmes neurons. *Abbreviations: HF, high-frequency; LF, low-frequency; NVmes, mesencephalic trigeminal nucleus*.

**TABLE 2.**
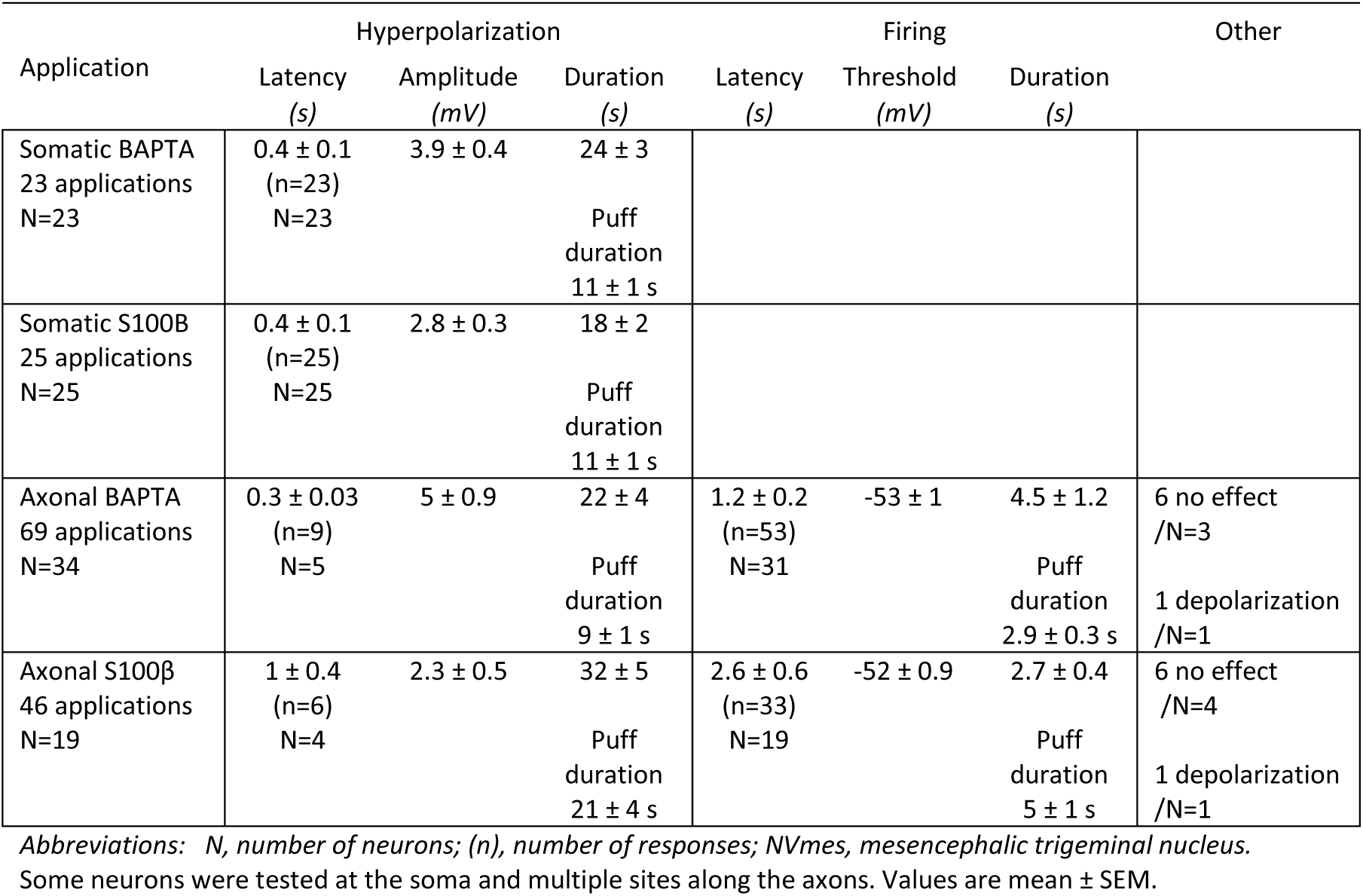
Effects of BAPTA and 100β applications on NVmes neurons.

| Application | Hyperpolarization |  |  | Firing |  |  | Other |
| --- | --- | --- | --- | --- | --- | --- | --- |
|  | Latency<br>(s) | Amplitude<br>(mV) | Duration<br>(s) | Latency<br>(s) | Threshold<br>(mV) | Duration<br>(s) |  |
| Somatic BAPTA<br>23 applications<br>N=23 | $0.4 \pm 0.1$<br>(n=23)<br>N=23 | $3.9 \pm 0.4$ | $24 \pm 3$<br>Puff<br>duration<br>$11 \pm 1$ s | | | | |
| Somatic S100β<br>25 applications<br>N=25 | $0.4 \pm 0.1$<br>(n=25)<br>N=25 | $2.8 \pm 0.3$ | $18 \pm 2$<br>Puff<br>duration<br>$11 \pm 1$ s | | | | |
| Axonal BAPTA<br>69 applications<br>N=34 | $0.3 \pm 0.03$<br>(n=9)<br>N=5 | $5 \pm 0.9$ | $22 \pm 4$<br>Puff<br>duration<br>$9 \pm 1$ s | $1.2 \pm 0.2$<br>(n=53)<br>N=31 | $-53 \pm 1$ | $4.5 \pm 1.2$<br>Puff<br>duration<br>$2.9 \pm 0.3$ s | 6 no effect<br>/N=3<br>1 depolarization<br>/N=1 |
| Axonal S100β<br>46 applications<br>N=19 | $1 \pm 0.4$<br>(n=6)<br>N=4 | $2.3 \pm 0.5$ | $32 \pm 5$<br>Puff<br>duration<br>$21 \pm 4$ s | $2.6 \pm 0.6$<br>(n=33)<br>N=19 | $-52 \pm 0.9$ | $2.7 \pm 0.4$<br>Puff<br>duration<br>$5 \pm 1$ s | 6 no effect<br>/N=4<br>1 depolarization<br>/N=1 |

Abbreviations: N, number of neurons; (n), number of responses; NVmes, mesencephalic trigeminal nucleus. Some neurons were tested at the soma and multiple sites along the axons. Values are mean ± SEM.

### Axonal applications of the astrocytic calcium-binding protein, S100β, cause firing in NVmes neurons

Given its ability to bind Ca^2+^, S100β, a protein synthesized and released by astrocytes, (26), is the physiological counterpart to BAPTA. The physiological roles of this protein are not all yet determined but our recent work suggests that one of its overlooked effects could be modulation of neuronal excitability through modulation of [Ca^2+^]_e_ (20, 21). We tested the effects of locally applied S100β near the soma and/or the stem axon of 31 NVmes neurons from 21 mice. S100β applied near the soma caused a hyperpolarization that outlasted the application by about 7s in all neurons tested (**Table 2**, **Fig 2A**, right trace). Forty-six applications of S100β were made near the axons of 19 NVmes neurons. As with BAPTA axonal applications, the majority of these applications (33/46) elicited firing (Table 2, **Fig 2B**, second, third, and fourth right traces), while a few (6/46) caused a membrane hyperpolarization (**Table 2**, **Fig 2B**, top right trace), a depolarization (1/46) or no effects (6/46). The elicited firing was most often preceded by either a depolarizing plateau (1.3 ± 0.1 mV, latency: 0.4 ± 0.1 s, n=17, **Fig 2B**, second right trace) or a hyperpolarization (2.8 ± 0.6 mV, latency: 0.5 ± 0.2 s, n=10, **Fig 2B**, third right trace). In the remaining 6 cases, firing emerged directly from the resting potential without prior membrane potential change at a latency of 2.2 ± 1.3 s (**Fig 2B**, bottom right trace). As with BAPTA, different firing patterns were observed with the most prevalent (9/33) being bursting (interburst frequency: 7.2 ± 1.4 Hz and an intra-burst frequency: 68 ± 6 Hz). Low (3.8 ± 1.4 Hz, n=8) and high (52 ± 10 Hz, n=5) frequency trains were also observed alone or with a mix of bursting (n=6 and 3 for low- and high-frequency trains, respectively). The remaining two applications elicited a single action potential at a latency of 9 and 16 s. The vertical bar charts in **Figure 3B** and **3C** illustrate the relative distribution of the different responses and the different firing patterns, respectively, observed in response to the S100β (purple bars) application near the soma or axonal process of the recorded neurons.

### Axonal applications of BAPTA or S100β increase NVmes neurons’ excitability by increasing SMOs and decreasing oscillation and firing threshold

Studies in biophysics have shown that extracellular Ca^2+^ ions exert a voltage-dependent partial block of Na^+^ channels and that removal of this block not only increases the amplitude of the current in single-channel measurements (19) but also shifts the activation gating range towards more hyperpolarized potentials (27). Thus, we first examined whether BAPTA and S100β altered the voltage-dependent firing and SMOs characteristics of NVmes neurons since both depend on Na^+^ channels. Given their similarity, BAPTA and S100β elicited firing responses were pooled together in this section for a total of 86 responses from 50 neurons. **Figure 3D** illustrates the distribution of these 86 firing responses among the 3 types of recorded NVmes neurons. Those responses were elicited at an average RMP of -56 ± 1 mV and their threshold for firing was -52 ± 1 mV (**Table 3**). Subthreshold membrane oscillations could be seen in 35 of the 86 firing responses from 28 neurons, 18 of which exhibited no SMOs before. In those responses, the SMOs were measured at an average RMP of -54 ± 1 mV, had an amplitude of 2.3 mV ± 0.1 mV, and a frequency of 71 ± 3 Hz (**Table 3**). A single response per neuron was considered for comparisons between controls and BAPTA or S100β induced effects (n=50). BAPTA and S100β triggered firing at more hyperpolarized potentials than did standard step current injections (**Fig 4A**; **Table 3 and Fig 4B**, empty and solid black bars, Wilcoxon Signed-rank test, *P* ˂ 0.001, n=50). The shift of firing threshold towards hyperpolarized potentials was also seen when using a ramp current injection protocol (**Fig 4C**, red arrows; **Table 3** and **Fig 4B**, empty and solid grey bars, Student paired *t*-test, *P* ˂ 0.001, n=7).

**Fig 4.**
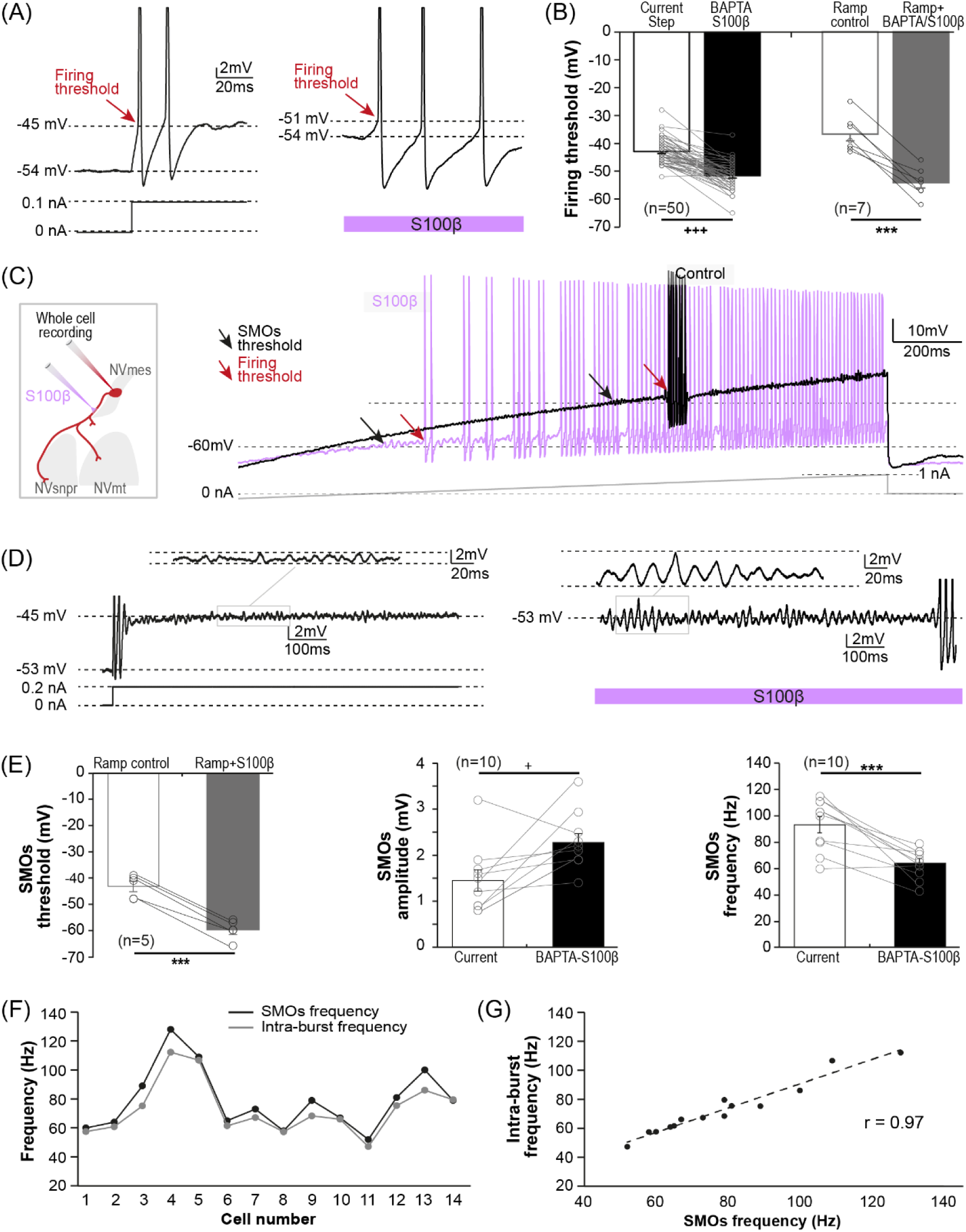
Lowering of SMOs and firing voltage thresholds of NVmes neurons by chelation of extracellular Ca^2+^ around their axonal processes. **(A):** Membrane voltage recordings of a NVmes neuron firing induced with depolarizing step current injection (**left** trace) or axonal application of S100β (**right** trace, as illustrated in the cartoon in **C**) showing the lowering of firing threshold with S100β in this neuron. **(B):** Bar chart of the firing voltage threshold of 50 NVmes neurons with step current injections (**empty black** bar, as illustrated in **A, left** trace) and with BAPTA and S100β axonal applications (**solid black** bar, as illustrated in **A, right** trace). The gray bars show the firing voltage threshold of 7 NVmes neurons with ramp current injection (as illustrated in **C**) in control condition (**empty grey** bar) and during BAPTA and S100β applications near the axonal process (**solid grey** bar) of the recorded neurons. **(C, left):** Cartoon illustrating the experimental set-up. **(C, right):** Recording of a NVmes neuron membrane potential while applying a ramp current injection (-1nA to 1nA, the hyperpolarizing part is truncated) in control condition (**black** trace) or during an axonal application of S100β (**mauve** trace). This condition triggered neuronal repetitive firing by lowering the SMOs (**black arrows**, control: -39 mV vs S100β: -59 mV) and firing (**red arrows**, control: -34 mV vs S100β: -57 mV) voltage thresholds compared to the control. (**D**): Membrane voltage recordings of a NVmes neuron SMOs induced with depolarizing step current injection (**left** trace) or axonal application of S100β (**right** trace, as illustrated in the cartoon in **B**) showing the increased amplitude of the SMOs with S100β in this neuron. **(E, left):** Bar chart of the SMOs voltage threshold of 5 NVmes neurons with ramp current injection in control condition (**empty grey** bar) and during S100β applications near the axonal process (**solid grey** bar) of the recorded neurons. **(E, middle):** Bar chart of the amplitude of the SMOs induced with depolarizing step current injection (**empty black** bar, as shown in **D, left**) or axonal application of BAPTA or S100β (**solid black** bar, as shown in **D, right**) in 10 NVmes neurons. **(E, right):** Bar chart of the frequency of the SMOs induced with depolarizing step current injection (**empty black** bar, as shown in **D, left**) or axonal application of BAPTA or S100β (**solid black** bar, as shown in **D, right**) in 10 NVmes neurons. **(F):** Plot of the intra-burst firing frequency (**grey** dots) and SMOs frequency (**black** dots) in the BAPTA- or S100β-induced burst-firing responses in 14 NVmes neurons showing how they parallel each other. (**G**): Scatterplot of the intra-burst firing frequency and the SMOs frequency in the BAPTA- or S100β-induced burst-firing responses showing a positive relationship between both variables. A linear regression (dotted line) between both variables shows that they are significantly correlated (r = 0.97, *P* < 0.001). *Abbreviations: NVmes, mesencephalic trigeminal nucleus; NVmt, trigeminal motor nucleus; NVsnpr, trigeminal main sensory nucleus.* ^+^ *P* ˂0.05, Wilcoxon Signed-rank test; ^+++^ *P* ˂ 0.001, Wilcoxon Signed-rank test; *** *P* ˂ 0.001, Student paired t-test.

**TABLE 3.**
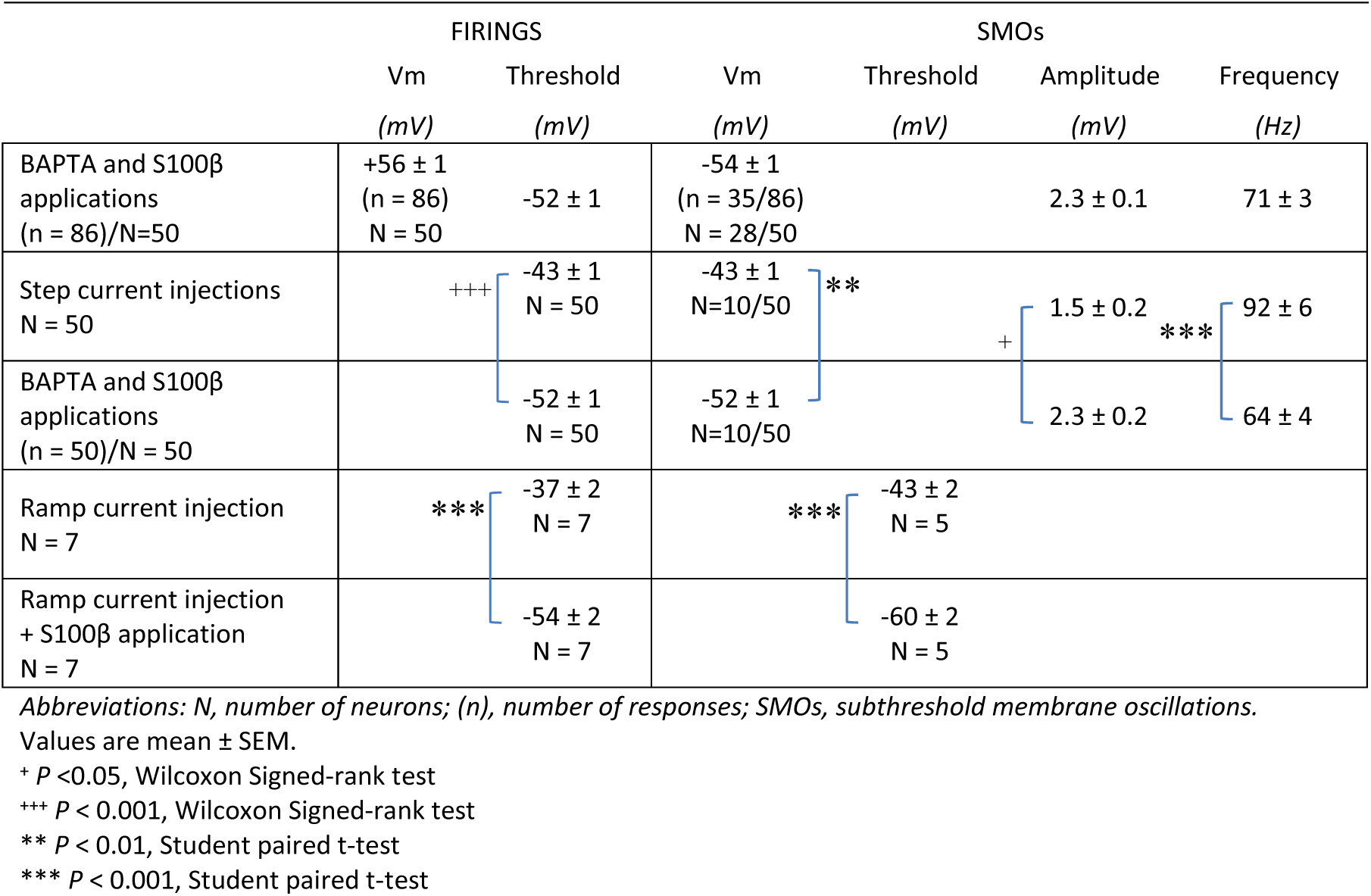
Effects of BAPTA and S100β axonal applications on firing threshold and SMOs characteristics.

Similar observations were made when comparing SMOs that appear intermingled with the firing during step current injection (n=10) or ramp current injection protocols (n=5) to those elicited during BAPTA and S100β-induced firings. The latter appeared, during ramp current injection, at potentials 10-16mV more hyperpolarized than the former (**Fig 4C**, black arrows; **Table 3 and Fig 4E**, left vertical bars chart, Student paired *t*-test, *P* ˂ 0.001, n=5). They also had a larger amplitude (**Fig 4D**; **Table 3 and Fig 4E**, middle vertical bars chart, Wilcoxon Signed-rank test, *P* = 0.02, n=10) and a lower frequency (**Table 3 and Fig 4E**, right vertical bars chart, Student paired *t*-test, P ˂ 0.001, n=10) than the SMOs induced by depolarizing steps current injection. When measured in the bursting responses (n=15/20 bursting responses) of 15 neurons, the SMOs frequency (**black** dots in the graph in **Fig 4F**) in each cell paralleled the intra-burst firing frequency (**grey** dots in the graph in **Fig 4F**) and both were significantly correlated (**Fig 4G**, linear regression, r = 0.97, P ˂ 0.001).

### *I*NaP is responsible for BAPTA- or S100β-induced firing

SMOs and bursting in NVmes neurons rely on *I*_NaP_ (12) which in these neurons is associated partly with the Na_V_1.6 isoform since it is significantly reduced, but not completely abolished in Na_V_1.6 null mice (25). To ascertain the involvement of *I*_NaP_ and Na_V_1.6 containing channels in the firing-inducing action of BAPTA and S100β, we tested the effects of Riluzole (a blocker of *I*_NaP_) and 4,9-anhydro-TTX (a blocker of sodium channels containing the Na_V_1.6 α subunit (28)). The BAPTA-induced firing was abolished by bath-application of either Riluzole (20 μM; not shown, n=2/2,) or 4,9-anhydro-TTX (0.1 µM; **Fig 5A**, n=7/7). In two additional cells, the S100β-induced firing was also abolished by bath applications of 4,9-anhydro-TTX. However, the SMOs accompanying 7 of these firing responses were blocked in 3 cases and only slightly reduced in amplitude in the 4 remaining cases (2.2 ± 0.6 mV vs 1.5 ± 0.6 mV; *P* = 0.109, Wilcoxon Signed-rank test). The input resistance and firing threshold of the recorded neurons were unchanged by 4,9-anhydro-TTX. However, the resting membrane potential was slightly but significantly more hyperpolarized upon 4,9-anhydro-TTX application (-56 ± 3 mV vs -58 ± 4 mV, *P* = 0.016, Student paired *t-*test), suggesting that *I*_NaP_ may contribute to the resting membrane potential in these cells.

**Fig 5.**
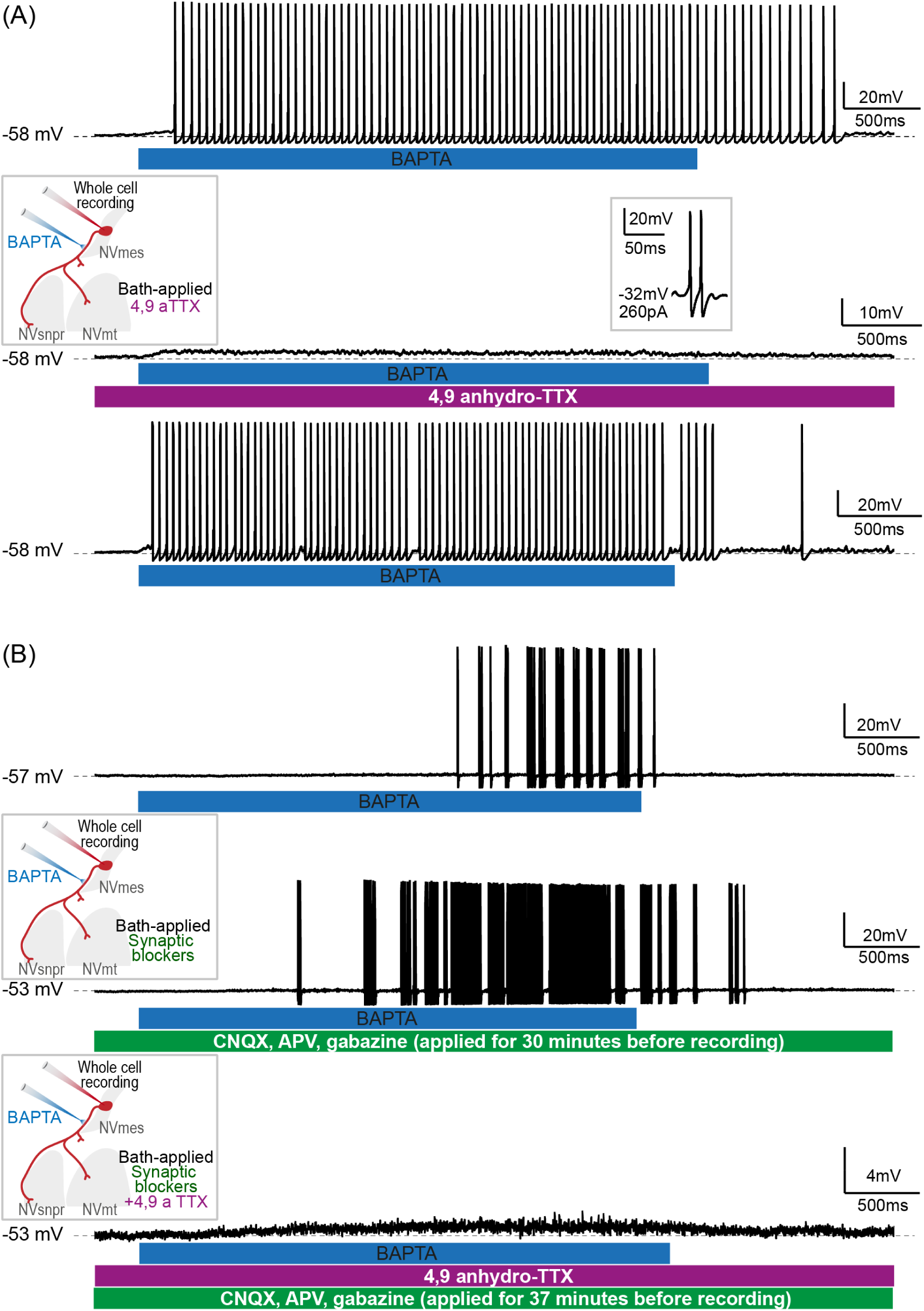
Axonal BAPTA- and S100β-induced firing in NVmes neuron persists in presence of synaptic blockers and appears to depend solely on I_NaP_ and the activity of Na_V_1.6 channels. **(A)**: Sustained firing recorded in a NVmes neuron following local application of BAPTA on its axonal process **(Top)** is abolished in presence of 4.9-anhydroTTX, a highly specific Na_V_1.6 blocker **(Middle)** and recovered after 40 min of washout of the 4,9-anhydroTTX (**Bottom**). The experimental setup is illustrated in the **Left inset,** while the **right inset** shows that the neuron is still capable of firing with membrane depolarization by current injection in presence of 4,9-anhydroTTX. **(B)**: Repetitive bursting recorded in a NVmes neuron following local application of BAPTA on its axonal process **(Top)** persists after 30 minutes of bath application of glutamatergic and GABAergic blockers (respectively CNQX, APV and gabazine; **Middle**), but is abolished by addition of 4,9-anhydroTTX to the perfusion (**Bottom)**. **Left insets**: illustration of the experimental setup. *Abbreviations: NVmes, mesencephalic trigeminal nucleus; NVmt, trigeminal motor nucleus; NVsnpr, trigeminal main sensory nucleus*.

To ensure that the firing induced in NVmes neurons by axonal application of BAPTA or S100β did not involve some indirect activation of synaptic transmission, which may have been masked by the potent effect of the *I*_NaP_ blockers, we tested the effect of a cocktail of antagonists for the AMPA/kainate (CNQX, 10 µm), NMDA (AP5, 26 µm), and GABAA (gabazine, 20 µm) receptors. Bath application of the synaptic blockers cocktail did not abolish the firing triggered by local application of BAPTA (**Fig 5B**, middle trace) or S100β along the axonal process of 7 tested neurons (BAPTA, n=4, S100β, n=3). Further application of 4,9-anhydro-TTX in addition to the synaptic blocker, in 3 out of these 7 pharmacological tests, successfully blocked the BAPTA, (n=1, **Fig 5B**, bottom trace) or S100β-induced (n=2) firing.

To further ascertain the requirement of Na_V_1.6 channels involvement in BAPTA or S100β-induced firing, both drugs were applied in the vicinity of NVmes neurons in Na_V_1.6 knockout mice. These mice express a mutation identified as a small LINE element insertion into exon 2 of the Scn8a gene which as a result encodes for a very short inactive protein (29). The 15 neurons recorded from 5 Na_V_1.6 knockout mice differed from those of WT/GFAP-ChR2 mice only for their firing threshold (-37 ± 2 mV; Kruskall Wallis test with Bonferroni correction, *P* = 0.03; **Table 1**). All showed an outward rectification during depolarization and an inward rectification during hyperpolarization (**Fig 6A**, right and left, red straight line: linear fitting to the non-rectifying portion of the I-V curve) producing a prominent sag upon membrane hyperpolarization (**Fig 6A**, arrow). All but one (14/15) showed, as the spike-adaptative subtype neurons, accommodation of firing upon membrane depolarization, with 12 of them discharging only a single action potential, even with long-duration pulses (up to 1000 ms). Local applications of BAPTA were tested in 10 NVmes neurons with 5 somatic applications in 5 cells and 11 axonal applications in 7 cells. Only two of the somatic applications produced an effect which was a membrane hyperpolarization (of 0.7 and 1.2 mV at a latency of 0.4 and 0.6 s; **Fig 6B**, top trace), while axonal applications caused membrane hyperpolarization (-1.3 mV) once, depolarization in 3 cases (1.1 ± 0.2 mV), firing once (in one of the spike-adaptative neurons), and no response in the 6 remaining cases (**Fig 6B**, bottom trace).

**Fig 6.**
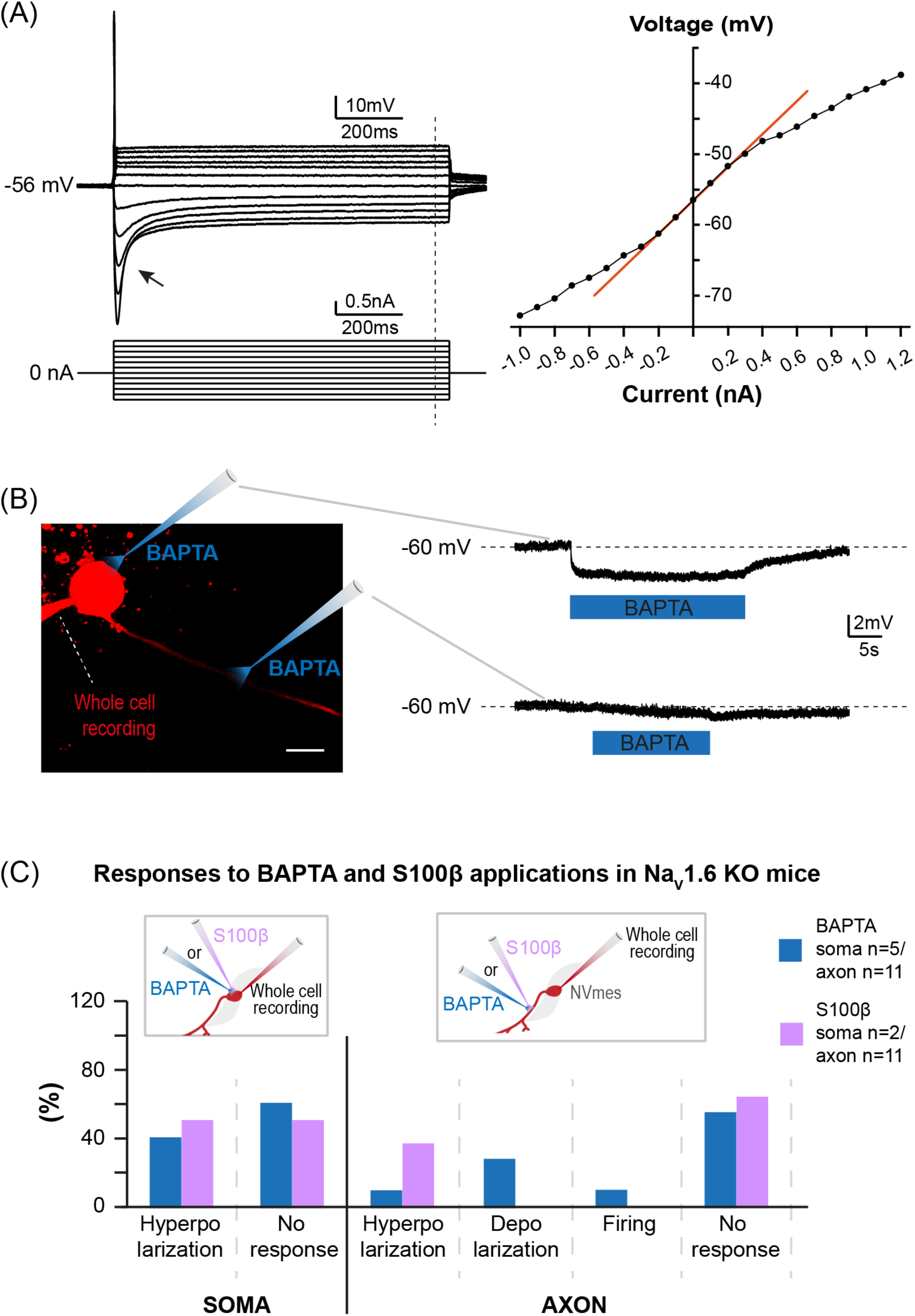
Nav1.6 KO NVmes neurons do not respond to axonal applications of BAPTA and S100β. **(A) Left**: Membrane responses of a Na_V_1.6-knock-out NVmes neuron (**top** traces) to injections of hyperpolarizing and depolarizing current pulses (**bottom** traces). The arrow points to the sag. The vertical dotted line indicates the position of the membrane voltage and the current injection lecture for the I-V curve. **Right**: I-V curve showing the inward and outward membrane rectification of a spike-adaptative Na_V_1.6-knock-out NVmes neuron in response to hyperpolarizing and depolarizing injected current, respectively. The red line is the linear fitting to the non-rectifying portion of the I-V curve. **(B) Left**: Photomicrography of a Na_V_1.6-knock-out NVmes neuron filled with intrapipette Alexa 594 with the BAPTA containing micropipette drawn to indicate the positions of the local applications, scale bar 20µm). **Right**: Membrane responses of the recorded neuron to somatic (**top** trace) and axonal applications (**bottom** trace). **(C):** Bar chart quantifying the response profiles of NVmes neurons to somatic or axonal applications of BAPTA (**blue**), or of S100β (**purple**) in Na_V_1.6-KO mice.

Local applications of S100β were tested on 4 NVmes neurons from 2 Na_V_1.6 knockout mice, with 2 somatic and 11 axonal applications. On the 2 somatic applications, only one produced a hyperpolarization (2.5 mV at a latency of 0.1 s), while the other had no effect. Seven of the axonal applications had no effect as well, while 4 caused a small hyperpolarization of 1.4 ± 0.1 mV at a latency of 0.5 ± 0.3 s. **Figure 6C** illustrates the relative distribution of the responses elicited by somatic and axonal applications of BAPTA or S100β (blue and purple bars, respectively) on NVmes neurons from Na_V_1.6 knockout mice.

### A precise axonal subregion is involved in the Nav1.6-dependent firing

Knowing that the BAPTA and S100β-induced firing is *I*_NaP_-dependent and that the Na_V_1.6 channels that support *I*_NaP_ are enriched in the AIS and the nodes of Ranvier, we tried to narrow down the precise locations at which local decrease of [Ca^2+^]_e_ were more potent to elicit firing. Given the simple morphology of these neurons, this was assessed by making very small and controlled applications of BAPTA or S100β at several locations (1 to 9) along the length of the axonal process of 34 neurons that were filled with Alexa fluor 594 or 488 through the patch pipette (as shown in **Fig 7A**). Great care was taken so that the BAPTA or S100β applied along the single process emerging from the cell body (length 173 ± 34 µm) flew away and not towards the soma. For each of these cases, the exact pipettes position was recorded in bright-field images and drawn offline over the course of the axon (as shown in **Fig 7A**). In this example, the first application of BAPTA, targeting the soma, produced a hyperpolarizing response (**Fig 7A**, top trace) while positions 2 and 3 (at 57 and 106 µm from the soma, respectively) produced sustained firing (**Fig 7A**, 2^nd^ and 3^rd^ traces). BAPTA applications further along the axon had no effects (**Fig 7A**, bottom traces). A total of 86 applications of BAPTA and S100β were made along the axons of the 34 recorded neurons. The vertical bar chart in **Figure 7B** illustrates the distribution of responses evoked in relation to the location of the application. Hyperpolarizing responses (grey bars) were mostly elicited by applications less than 40 µm from the soma while firing responses (black bars) were predominantly produced by applications positioned between 40 and 100 µm from the soma. Among the firing responses, those that occurred concomitantly with a membrane potential hyperpolarization (**Fig 7C**, gray bars) were triggered by more proximal applications, while those accompanied by a depolarizing plateau (**Fig 7C**, black bars) were elicited by applications at any level of the axon.

**Fig 7.**
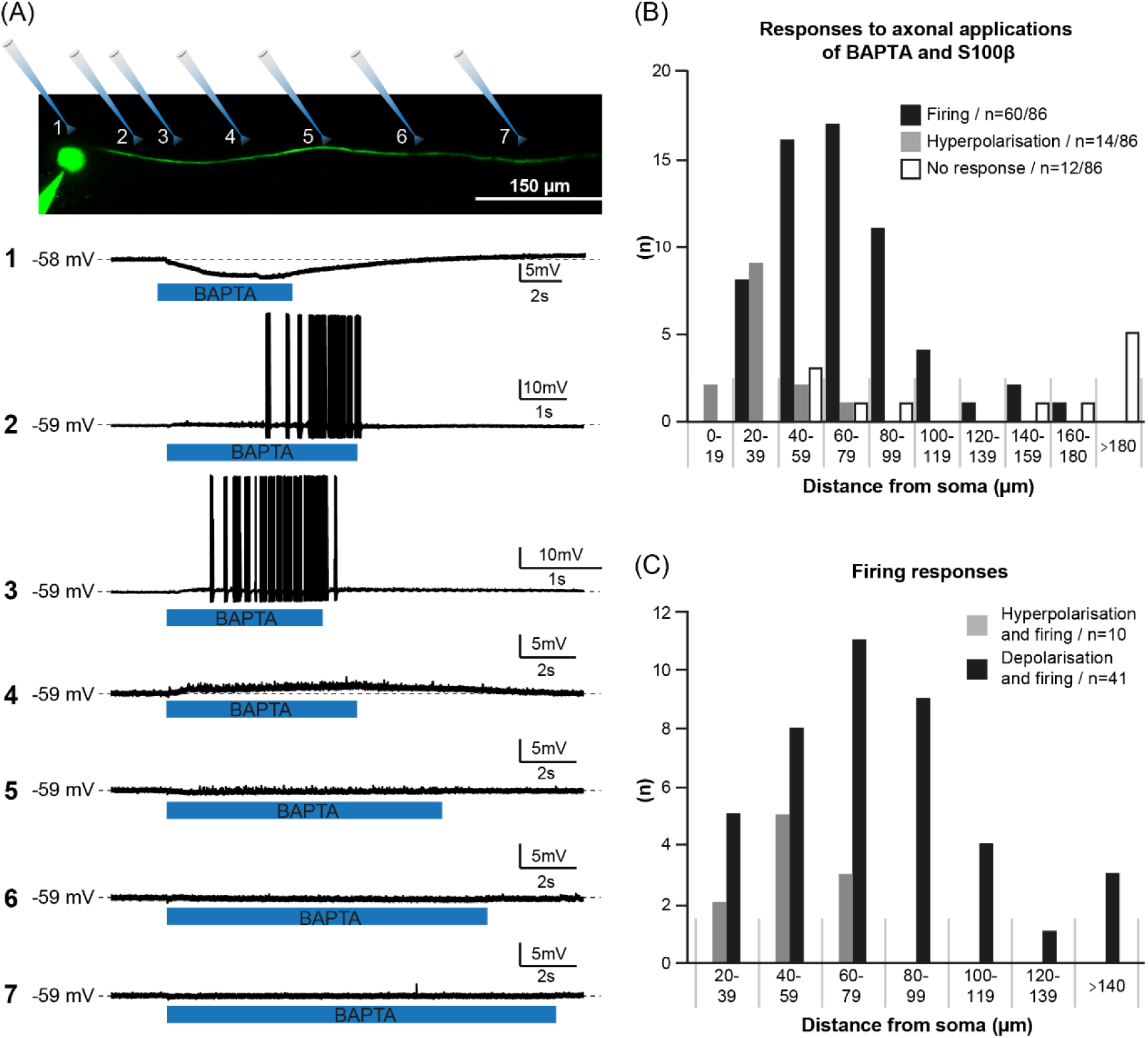
Chelation of extracellular Ca^2+^ around specific axonal subregions initiate NVmes neuronal bursting. **(A) Top:** Reconstructed image of the tested NVmes neuron filled with intrapipette Alexa 488, indicating all positions of BAPTA applications along the axon for the recorded membrane responses below. **Bottom 1-7:** Electrophysiological responses elicited by local BAPTA applications at the top-illustrated corresponding positions. Somatic application (position 1) of BAPTA induced a hyperpolarization, while bursting was initiated for the 2 and 3 micropipette positions, corresponding respectively to a 57 and 107 µm distance from the soma. Position 4 induced only a slight depolarization associated with membrane potential oscillations, and all further BAPTA applications had no effect. Initial membrane potentials and scale bars are indicated next to each trace. **(B):** Bar chart of the distribution of all neuronal responses to the axonal applications of BAPTA and S100β relatively to the distance from the soma of the local application. **(C):** Most firing responses were observed following a neuronal depolarization (**black** bars, n=41), but some neurons displayed firing following a hyperpolarization episode (**grey** bars, n=10).

### Co-localization of Nav1.6 channels and S100β-positive cells and processes around NVmes neurons fibers and somata

The above data reveal that BAPTA and S100β induce firing in a specific subregion of the axon of NVmes neurons, suggesting that there may be specific sites along the axons of these neurons that are more sensitive to extracellular calcium depletion. Thus, we hypothesized that Na_V_1.6 and the endogenous calcium chelator S100β may colocalize at these sensitive axonal locations. We used immunohistochemistry to examine the distribution of S100β-immunoreactive fibers along NVmes neurons’ cell bodies and axonal fibers. As many large diameter primary afferents, NVmes neurons are unipolar and can be positively identified using parvalbumin as in **Figure 8A** (n=3) which shows them to be surrounded by S100β-immunoreactive cell bodies (arrows) and fibers (arrowheads). Further, combined immunocytochemistry against Na_V_1.6 channels and S100β showed close apposition of S100β positive processes along Na_V_1.6 immunoreactive cell bodies and axons of the NVmes neurons in the WT mice (n=6; **Fig 8B)**. Only cell bodies were marked by immunocytochemistry against Na_V_1.6 in the Na_V_1.6 null mice (n=2), suggesting that if part of the mutated channel could still be targeted by the antibody, the mutation prevents its successful trafficking to its final position along the axonal process (**Fig 8C)**. Interestingly, NVmes somata are also S100β-immunoreactive. Similar observation was reported in the rat NVmes (30). A study using single RNA sequencing revealed that NVmes neurons express S100β (31), leading us to believe that the labeling observed in the cell bodies of these neurons is not artefactual.

**Fig 8.**
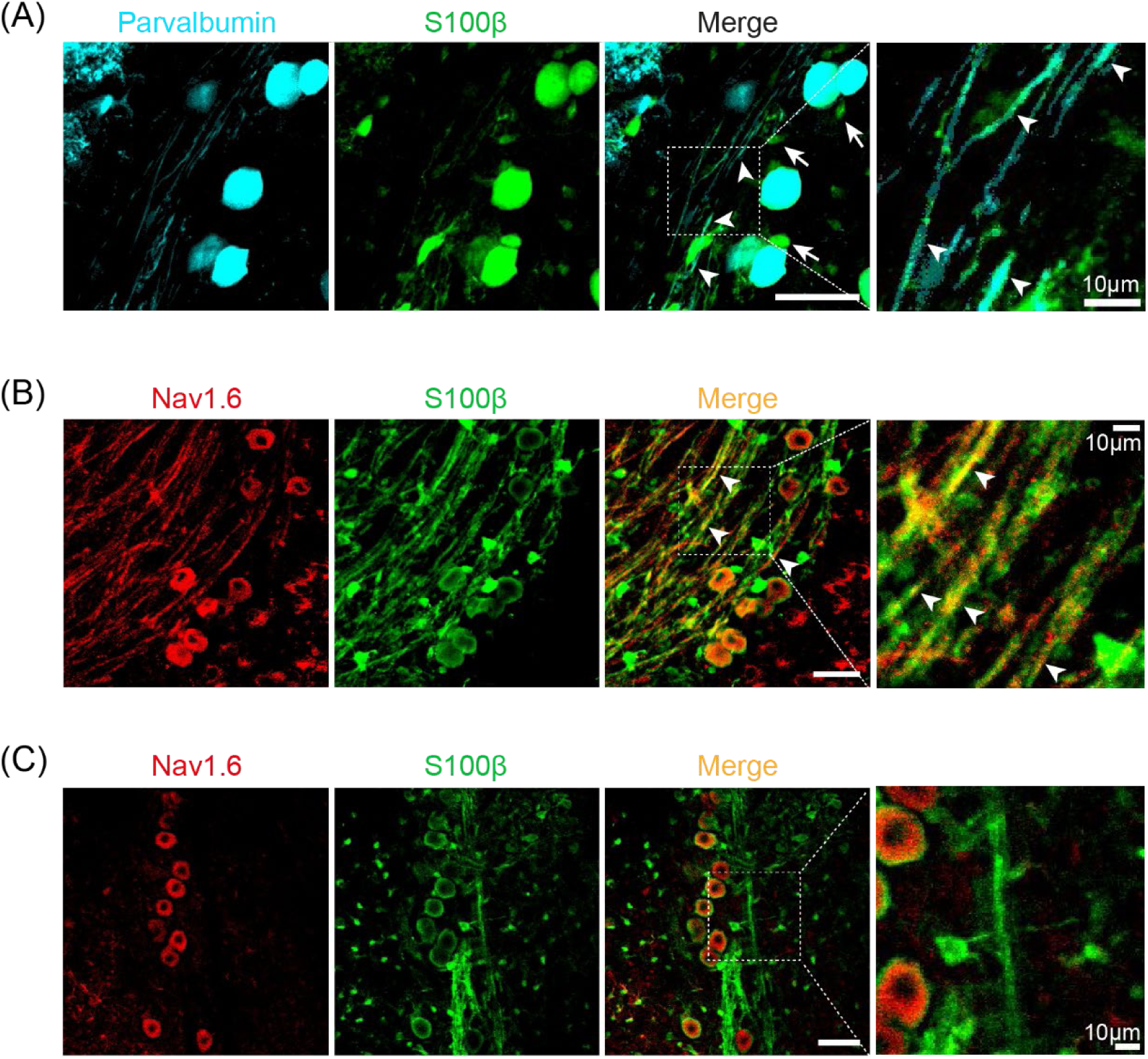
Close relationship between astrocytes and Na_V_1.6 immunopositive NVmes neurons. **(A):** Parvalbumin-positive NVmes neurons (cyan**, left**) cell bodies are surrounded by astrocytes (arrows) (S100β staining, **second,** green) which also contact NVmes axons (arrowheads). **(B, C):** Immunofluorescence staining of Nav1.6 (**left** panels, red), S100β (**second,** green) and superposition of Nav1.6 and S100β (**right,** yellow) in WT (**B**) and Nav1.6 KO (**C**) mice. Axonal colocalization of Nav1.6 and S100β indicating axonal apposition of astrocytic processes over Nav1.6 channels (arrowheads) is observed in WT, but not Nav1.6 KO mice. Scale bars 50 µm.

### Activation of astrocytes near NVmes neurons axons causes firing

Given that S100β is present in the vicinity of axonal Na_V_1.6 and because astrocyte activation/depolarization has been linked to astrocyte release of S100β (21, 32), we used mice expressing channel-rhodopsin 2 (ChR2) under the astrocytic GFAP promoter to examine the effects of stimulating surrounding astrocytes on NVmes neurons. The recorded neurons were filled with Alexa Fluor 594 through the patch pipette to visualize their cell body and axon and to allow manual delimitation of defined zones for optogenetic stimulation of neighboring astrocytes. Ninety neurons showing a round or ovoid cell body were recorded in the NVmes of 63 GFAP-ChR2 mice. A single process emerging from the cell body could be visualized in 73 of them (length 133 ± 9 µm). ROIs were drawn to stimulate the astrocytes around the soma only in 26 neurons (**Fig 9A**), around the soma and the axonal process in 10 neurons (**Fig 9B)**, and around the axonal process only in 67 neurons (**Fig 9C),** respectively. The parameters of the photostimulated areas were measured offline for the cases where images were recorded (**Table 4**). In some cells, more than one neuronal compartment was tested. The recorded neurons showed similar electrophysiological properties to the neurons recorded in wild-type mice (see **Table 1**).

**Fig 9.**
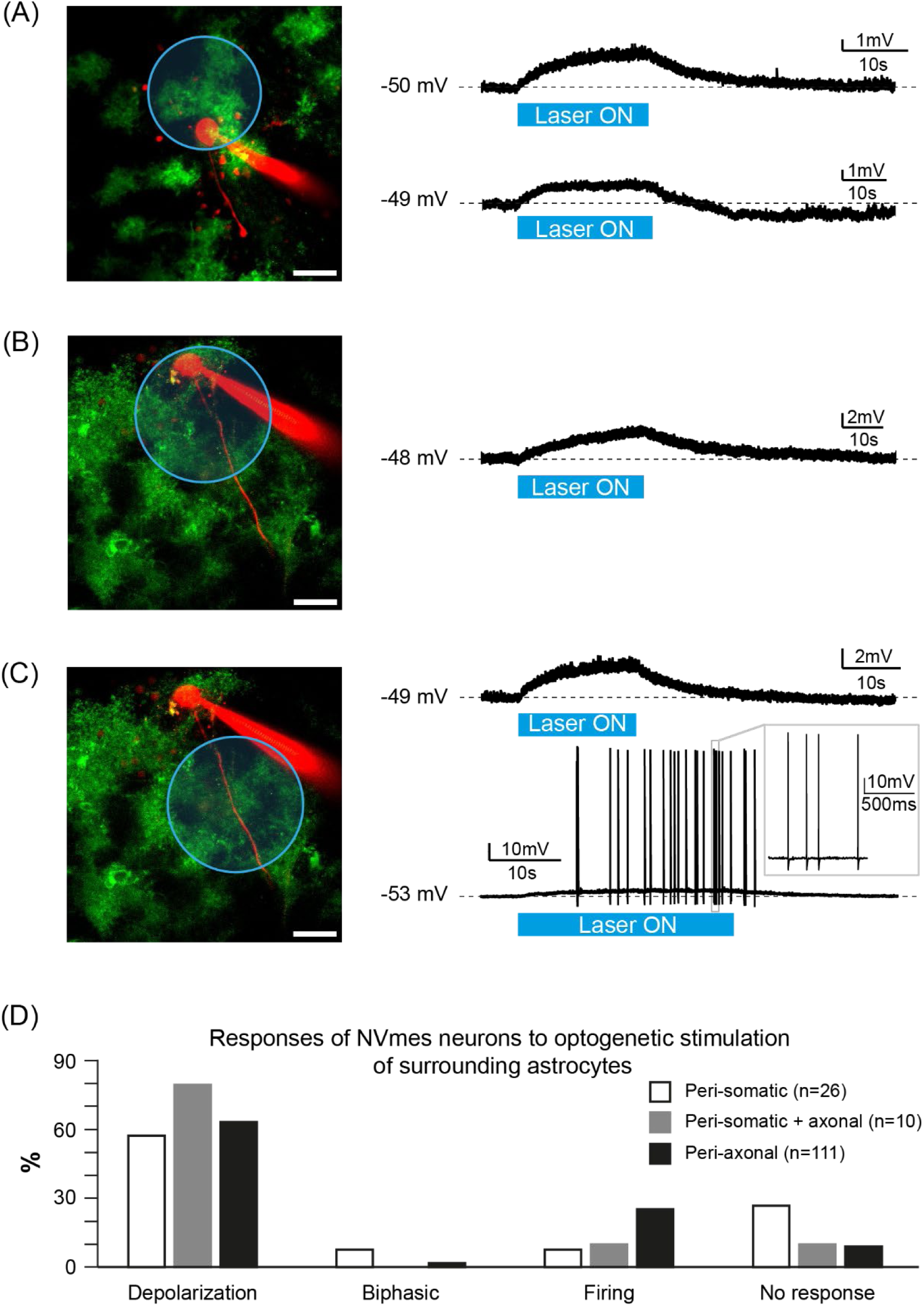
Optogenetic stimulation of peri-neuronal astrocytes in GFAP-ChR2-EYFP mice elicits a diversity of effects. **(A, B and C):** Photomicrographs of the area of optogenetic stimulation (**blue** circle) of peri-somatic astrocytes **(A)**, or both peri-somatic and peri-axonal astrocytes **(B)** or only peri-axonal astrocytes **(C)** (green astrocytes, red through intrapipette Alexa-594 recorded NVmes neuron), scale bar 50 µm. **(A:)** Example of a long-lasting depolarization response (**top** trace) and of a biphasic response consisting of a depolarization followed by a hyperpolarization (**bottom** trace) induced by the optogenetic stimulation of peri-somatic astrocytes. **(B):** Example of a long-lasting depolarization response induced by the optogenetic stimulation of both peri-somatic and peri-axonal astrocytes. **(C):** Example of a long-lasting depolarization response (**top** trace) and of repetitive firing (**bottom** trace) induced by the optogenetic stimulation of peri-axonal astrocytes. **(D):** Bar chart of the responses of NVmes neurons to optogenetic stimulation of peri-somatic (**white**), peri-somatic and peri-axonal (**grey**) and only peri-axonal (**black**) astrocytes.

**TABLE 4.** Stimulated areas.

| Surface | Distance<br>From the soma | Length along<br>the axon |
| --- | --- | --- |

| | ($\mu\text{m}^2$) | ($\mu\text{m}$) | ($\mu\text{m}$) |
| --- | --- | --- | --- |
| **Peri-Somatic** | 8240 ± 2227<br>(n = 20) | Soma included | 0 |
| **Peri-Somatic + Axonal** | 23439 ± 6940<br>(n = 6) | Soma included | 89 ± 21 |
| **Peri-Axonal** | 8589 ± 585<br>(n = 93) | 52 ± 4 | 65 ± 4 |

Values are mean ± SEM. (n = number of images analyzed)

#### Activation of the peri-somatic astrocytes

Optogenetic stimulation of astrocytes in areas encompassing the soma but excluding the axon of the recorded neuron generally led to a small depolarization (**Table 5**, **Fig 9A**, top trace; n=15) or no effect (n=7). Other responses observed included biphasic responses consisting of small long-lasting depolarization followed by a hyperpolarization (**Fig 9A**, second trace; n=2), or an increase in firing frequency in a cell that was spontaneously firing (not shown). Different stimulation durations were tested (from 10 to 40 s) to see if firing could be elicited, but even though the observed depolarizations outlasted the stimulation period, firing could not be elicited except for the cell that was spontaneously active, and one other cell where a single spike was produced. The size of the stimulated area was measured offline in 20 of the 26 cases where images were recorded (**Table 4**). The vertical bar chart in **Figure 9D** illustrates the relative distribution of the different responses elicited by the optogenetic stimulation of the astrocytes surrounding the cell body (empty bars) of the patched neurons.

**TABLE 5.** Effects of optogenetic stimulation of astrocytes on NVmes neurons.

|  |  | Depolarization |  |  | Firing |  | Other |  |
| --- | --- | --- | --- | --- | --- | --- | --- | --- |
|  |  | Latency | Amplitude | Duration | Latency | Duration | Biphasic | No effect |
|  |  | (s) | (mV) | (s) | (s) | (s) |  |  |
| Peri-Somatic (10-40 s)<br>(n = 26)<br>N = 26 | At RMP | 0.4 ± 0.1<br>(n=15)<br>N = 15 | 2 ± 0.2 | 17-77 | 17<br>2.2<br>(n = 2)<br>N = 2 | 0.03<br>28 | (n = 2)<br>N = 2<br>Lat.3.2<br>and 0.9 s | (n = 7)<br>N = 7 |
| Peri-Somatic + Axonal (30 s)<br>(n = 10)<br>N = 10 | At RMP | 0.6 ± 0.1<br>(n = 8)<br>N = 8 | 1.7 ± 0.4 | 50 ± 7 | 14<br>(n = 1)<br>N = 1 | 14 |  | (n = 1)<br>N = 1 |
| Peri-Axonal (5-30 s)<br>(n = 111)<br>N = 67 | At RMP<br>(n = 89)<br>N = 64 | 0.5 ± 0.1<br>(n = 67)<br>N = 51 | 1.7 ± 0.1 | 14-114 | 6.5 ± 1.1<br>(n = 11)<br>N = 10 | 0.03-112 | (n = 2)<br>N = 2<br>Lat. 0.3<br>and 0.7 s | (n = 9)<br>N = 7 |
|  | With depolarizing current<br>(n = 22)<br>N = 22 | 0.7 ± 0.4<sup>a</sup><br>(n = 4)<br>N = 4 | 2.5 ± 1<sup>b</sup> | 21-73<sup>c</sup> | 5.1 ± 0.5<br>(n = 17)<br>N = 17 | 47 ± 11 |  | (n = 1)<br>N = 1 |

Abbreviations: Lat., latency; N, number of neurons; (n), number of stimulation sites; NVmes, mesencephalic trigeminal nucleus; RMP, resting membrane potential.
Values are mean ± SEM for data that can be averaged; exact values for n < 3; or intervals for duration data acquired with different stimulation durations.
<sup>a</sup>; <sup>b</sup>; <sup>c</sup>: values not different from control for the 4 depolarizations tested at RMP: lat, 0.5 ± 0.2, Paired t-test, *P* = 0.422; amplitude, 2.8 ± 1.9, Paired t-test, *P* = 0.281; duration, 46 ± 19, Paired t-test, *P* = 0.91, respectively.

#### Activation of the astrocytes surrounding the cell body and the axonal process

In 10 cells, stimulation of larger areas that included the soma and the proximal part of the axon also produced long-lasting depolarizations in 8 cases (**Table 5**, **Fig 9B**). A volley of 14 action potentials (not shown) was elicited in one case, while no detectable response could be seen in the last tested case. The size of the photostimulated area and the length of exposed axon were measured offline in 6 of the 10 cases where images were recorded (**Table 4**). The vertical bars chart in **Figure 9D** illustrates the relative distribution of the different responses elicited by the optogenetic stimulation of the astrocytes surrounding the cell body and the proximal axon (grey bars) of the patched neurons.

#### Activation of the peri-axonal astrocytes

In 67 of the 90 patched neurons, an ROI drawn for photostimulating the astrocytes excluded the cell body and was restricted to part of the axonal process and the surrounding astrocytes. In some cells, more than one site was tested along the length of the axon and some sites were counted more than once because they were tested with different durations and/or with/without current injection for a total of 111 sites of stimulation (**Table 5**). The size of the photostimulated area, its distance from the soma, and the length of the exposed axon were measured offline in 93 of the 111 cases where images were recorded (**Table 4**). As with peri-somatic stimulation, when tested at the RMP, peri-axonal astrocytic optogenetic stimulation produced a long-lasting depolarization (**Table 5**; **Fig. 9C**, top trace) that outlasted the optogenetic pulse duration for 67 stimulation sites.

Biphasic responses were elicited by stimulating 2 other sites (**Table 5**, not shown), and firing overriding long-lasting plateaux (**Fig 9C**, bottom trace) were seen in 11 other neurons (2 of which were spontaneously firing). In the 9 remaining cases, the optogenetic stimulation caused no detectable response.

To facilitate propagation of the stimulation-evoked action potentials into the cell body of the recorded neuron, we tried pairing the optogenetic stimulation of the astrocytes surrounding the axon to membrane depolarization, using current injection, in 22 neurons; 18 of which had been tested at their RMP (-53 ± 1). One neuron did not respond to the optogenetic stimulation while the 17 remaining ones responded with a subthreshold long-lasting depolarization. Injection of depolarizing current (140 ± 16 pA, n=22) brought the membrane potential to -44 ± 1 mV (n=22). In 4 of these, prior depolarization did not change significantly the amplitude (Paired *t*-test, *P* = 0.281), the latency (Paired *t*-test, *P* = 0.422), or the duration (Paired *t*-test, *P* = 0.901) of the depolarization induced by astrocytic stimulation (**Table 5**). The non-responding neuron still failed to respond with prior depolarization. For 17 neurons, the pairing of the optogenetic stimulation of peri-axonal astrocytes with depolarizing current injection (which caused some firing by itself in 2/17 cases) triggered sustained firing overriding the long-lasting depolarizing plateau (as in **Fig 9C**, bottom trace). The vertical bars chart in **Figure 9D** illustrates the relative distribution of the different responses elicited by the optogenetic stimulation of the astrocytes surrounding the axon (black bars) of the patched neurons.

In summary, the optogenetic stimulation of peri-axonal astrocytes (tested in a total of 111 peri-axonal sites in 67 NVmes neurons) caused (n=24) or increased (n=4; Student paired *t*-test, *P* = 0.03) firing in 28 cases. As with the local applications of BAPTA and S100β, different firing patterns were observed in response to the stimulation of the peri-axonal astrocytes. The most prevalent pattern seen in 11 cases was bursting with the emission of a single burst 9 and 13 s after the start of the stimulation in 2 cases and repetitive bursting in the other 9 cases (**Fig 10A**, top trace; latency of 6 ± 1; duration of 16 ± 3 s; interburst frequency: 1 ± 0.3 Hz; intra-burst frequency: 65 ± 4 Hz). In one case, a single spike was elicited at a latency of 9.2 s while in 7 cases, low-frequency train of singlets was induced (**Fig 10A**, second trace; frequency 0.8 ± 0.2 Hz; latency: 6.4 ± 1 s; duration: 30 ± 14 s). High frequency (89 Hz) train intermingled with bursting (**Fig 10A**, third trace) was seen in a single cell, while 8 stimulations gave rise to a mix of low-frequency train of singlets and bursts (**Fig 10A**, bottom trace (latency: 3 ± 0.3 s; duration: 17 ± 5 s). The vertical bars chart in **Figure 10B** illustrates the relative distribution of the different firing patterns observed in response to the optogenetic stimulation of the peri-axonal astrocytes while **Figure 10C** illustrates the distribution of these 28 firing responses among the 3 types of recorded NVmes neurons.

**Fig 10.**
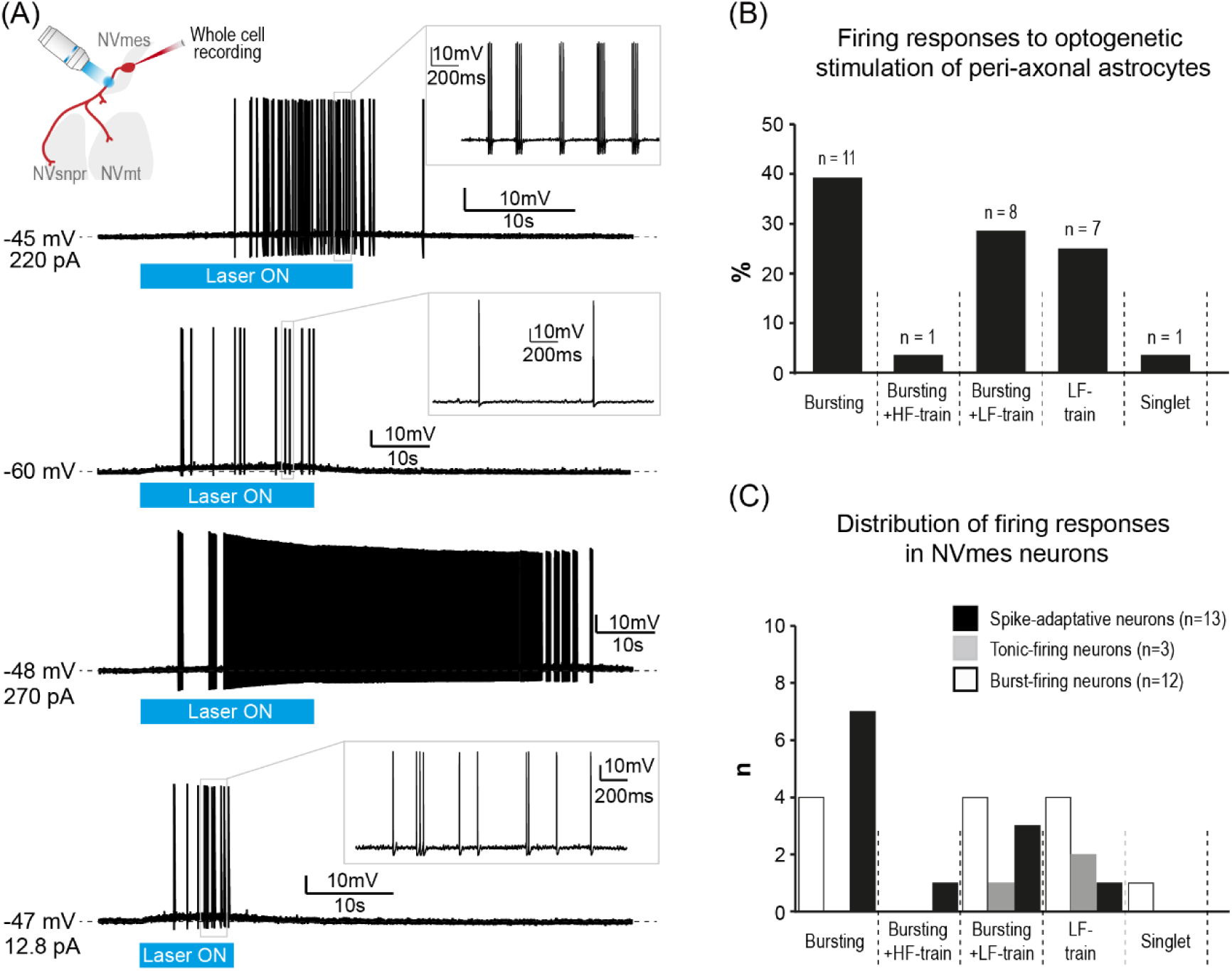
NVmes neurons firing responses to the optogenetic stimulation of peri-axonal astrocytes in GFAP-ChR2-EYFP mice. **(A):** The optogenetic stimulation (blue light, combination of 440 and 488 nm lasers, 15% laser power) of peri-axonal astrocytes (as illustrated by the cartoon) triggered diverse firing patterns in the recorded NVmes neurons including bursting (**top** trace), single spikes (**second** trace), bursting associated with single action potentials (**bottom** trace), and in one occurrence, action potentials train mixed with bursting (**third** trace). **(B):** Bar chart of the relative distribution of the firing responses (shown in **A**) elicited by peri-axonal astrocytes stimulations. **C):** Bar chart of the relative distribution of the neuronal firing responses elicited by peri-axonal astrocytes stimulations in the spike-adaptative (black bars), tonic-firing (grey bars), and burst-firing (empty bars) NVmes neurons. *Abbreviations: HF, high-frequency; LF, low-frequency; NVmes, mesencephalic trigeminal nucleus; NVmt, trigeminal motor nucleus; NVsnpr, trigeminal main sensory nucleus*.

Interestingly, in 19 cases, optogenetic stimulation of peri-axonal astrocytes caused an increase of the amplitude (1.7 ± 0.1 mV vs 2.4 ± 0.2 mV, Wilcoxon Signed-rank test, P ˂ 0.001) and a decrease of the frequency (74 ± 6 Hz vs 61 ± 4 Hz, Paired *t-*test, *P* = 0.041) of the SMOs. The optogenetic stimulation of peri-axonal astrocytes also triggered firing at slightly more hyperpolarized potentials than did standard step current injections (-45 ± 1 mV vs -47 ± 1 mV, Paired *t*-test, *P* = 0.002, n= 10) in 10 of the 11 neurons in which spikes were elicited at RMP.

### A precise axonal subregion is involved in the astrocytes-induced firing

We then sought to examine, using the images recorded for offline analysis, if, as with axonal applications of BAPTA and S100β, a specific subregion could be identified along the axonal process of the recorded neurons for the successful triggering of firing with optogenetic stimulation of peri-axonal astrocytes. We drew horizontal lines (blue, cyan, black, and gray lines in **Fig 11A-D**) along the axon of a schematic illustration of an NVmes sensory neuron (red pseudo-unipolar cells in **Fig 11A-D)** representing the length and distance from the soma of the portion of illuminated axon for each trial of optogenetic stimulation of peri-axonal astrocytes for 3 categories of evoked responses and for the cases where no responses could be detected. It appears that, without the aid of imposed depolarization, the firing responses (n=11, **Fig 11A**) were predominantly evoked by stimulation zones that cover the portion of the axon comprised between 25 and 80 µm from the soma. Membrane depolarization extends the length of the responsive axon proximally relatively to the soma (n=14, **Fig 11B**). Long-lasting depolarizations could be elicited at all levels of the axon length (n=58, **Fig 11 C**) while the cases of failures to respond are evenly distributed along the axonal length (n=10, **Fig 11D**). Those observations are summarized in the bar chart in **Figure 11E** which represents the responses evoked by the optogenetic stimulation of peri-axonal astrocytes in relation to the position of the proximal extremity of the covered area of stimulation.

**Fig 11.**
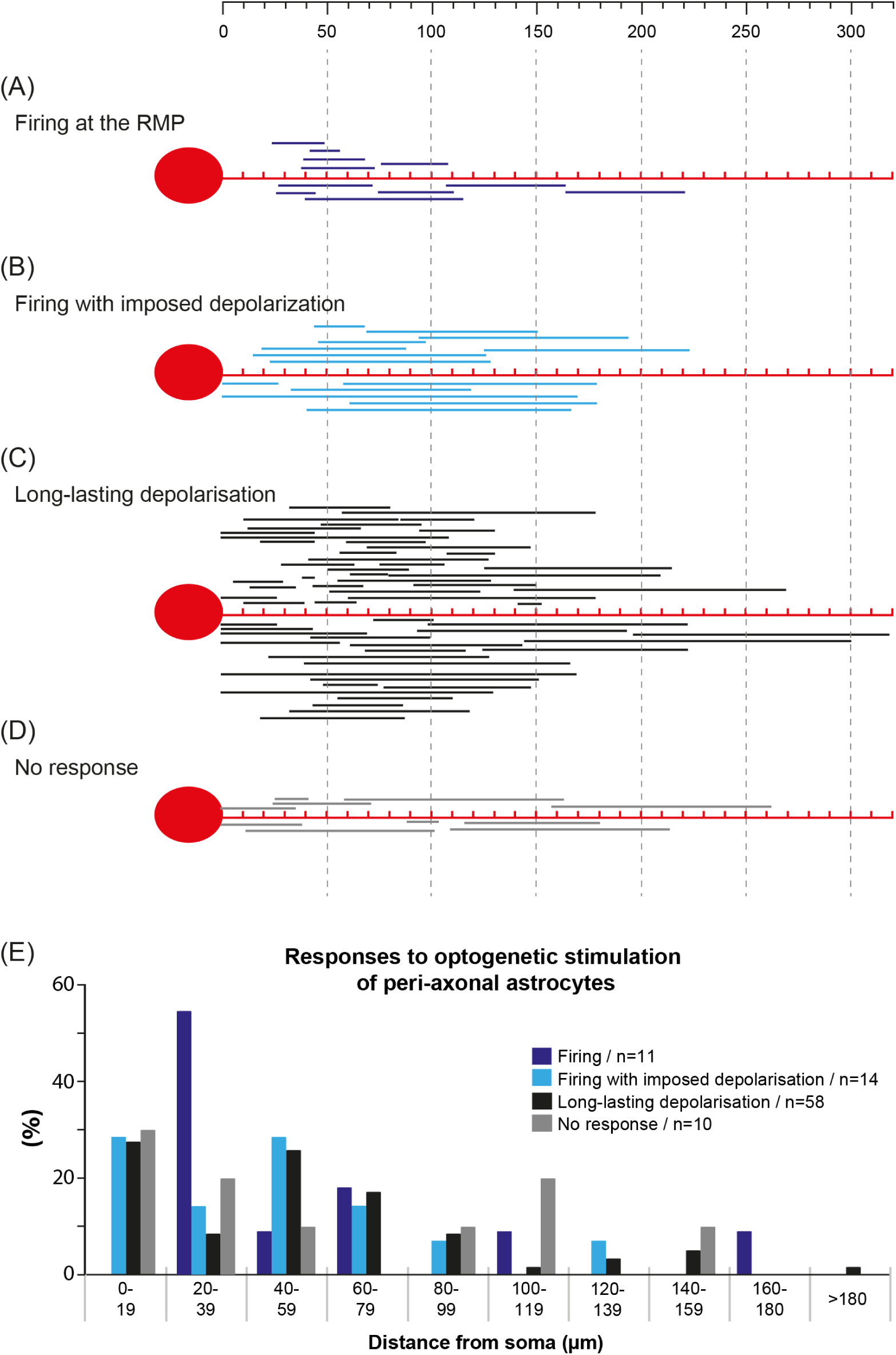
NVmes neuronal firing is initiated by optogenetic stimulation of peri-axonal astrocytes in GFAP-ChR2-EYFP mice surrounding specific axonal subregions. **(A-D):** Graphic summary of recorded responses following optogenetic stimulations of astrocytes surrounding NVmes axonal subregions in GFAP-ChR2-EYFP mice in relation to the position of the proximal extremity of the covered area of stimulation from 0µm up to 300µm of distance from the soma (top grid lines) for 93 of the 111 stimulation sites where images were recorded for offline analysis. All responses with the nominal distances of the proximal extremity of the covered area from the soma are plotted in **(E)**. Following these optogenetic stimulations, we observed firing at the resting membrane potential for 11 stimulation sites (**A; E** purple bars) and firing following an imposed depolarization for 14 stimulation sites (**B; E** blue bars). The most common response was a long-lasting depolarization elicited by 58 stimulation sites (**C; E** black bars), with 10 sites producing no response (**D; E** grey bars).

### The astrocytes-induced firing depends on S100β and Nav1.6 channels

To elucidate whether release of endogenous S100β in the extracellular space was involved in the observed effects, we tested whether local applications of an anti-S100β antibody (40–80 µg/ml) affected firing induced by optogenetic stimulation of neighboring astrocytes. **Figure 12A** (**top** trace) illustrates the response evoked in the patched neuron shown in the image by the optogenetic stimulation of the astrocytes (arrowheads) enclosed in the delineated zone (blue circle) around its axon. Local application of the S100β-subunit monoclonal antibody completely abolished the firing (**Fig 12A. middle** trace; n=5 neurons, from 5 slices, from 5 animals) induced by the optogenetic stimulation which gradually recovered (**Fig 12A, bottom** trace) after washout of the antibody. However, the small-amplitude underlying depolarizations remained in the presence of the anti-S100β antibody in the 5 tested cases (**Fig 12A. middle,** top trace above the dotted baseline in inset).

**Fig 12.**
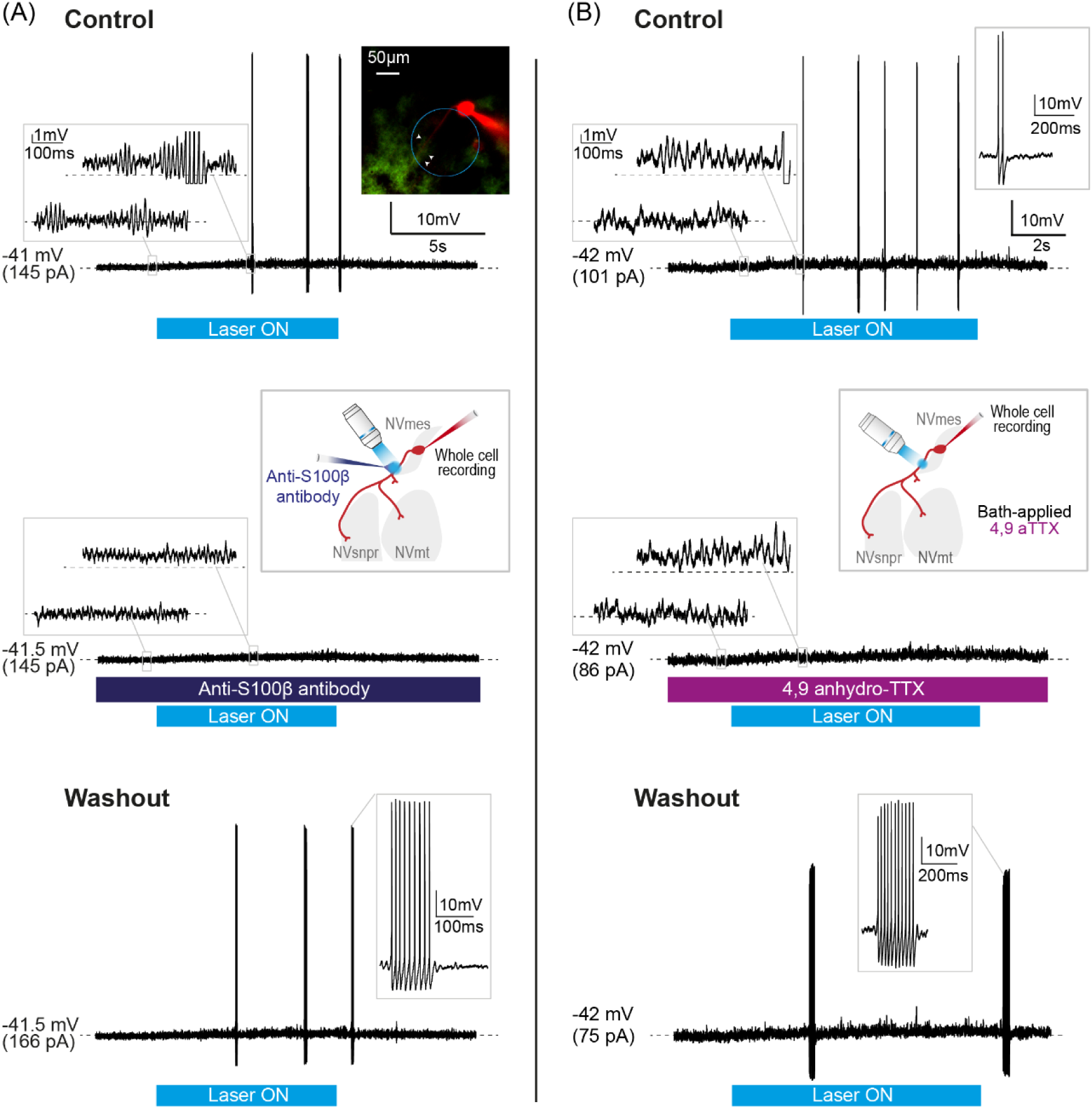
Firing triggered in NVmes neurons by optogenetic stimulation of peri-axonal astrocytes in GFAP-ChR2-EYFP mice is reversibly suppressed by the application of an S100β antibody and a Na_V_1.6 channels blocker. **(A):** Transient bursting **(Top** trace; Left Inset emphasizes the slight depolarisation underlying the firing) induced in an NVmes neuron by a 10 s optogenetic stimulation of its peri-axonal astrocytes (arrowheads in blue circle in photomicrograph) is abolished following local application of the anti-S100β antibody **(Middle;** Left inset emphasizes the remaining slight depolarisation. Right inset illustrates the experimental setup) and recovered after a 35-minute wash **(Bottom)**. **(B):** Doublets and singlets (**Top** trace; Left inset emphasizes the slight depolarisation underlying the firing) induced in an NVmes neuron by a 10 s optogenetic stimulation of its peri-axonal astrocytes are abolished following the addition of 4.9-anhydroTTX, a selective Nav1.6 antagonist, to the perfusion bath **(Middle;** Left inset emphasizes the remaining slight depolarisation. Right inset illustrates the experimental setup). **Bottom:** A bursting response was recovered after a prolonged wash. *Abbreviations: NVmes, mesencephalic trigeminal nucleus; NVmt, trigeminal motor nucleus; NVsnpr, trigeminal main sensory nucleus*.

To further determine if the effect of endogenous S100β also involved Na_V_1.6 channels, we tested the effect of bath-applications of 4,9-anhydro-TTX on the astrocytes-induced firing in 2 neurons (from 2 slices, from 2 animals). **Figure 12B** (**top** trace) illustrates the response evoked in one of these neurons by the optogenetic stimulation of the astrocytes surrounding its axon. Bath-application of 4,9-anhydro-TTX completely abolished the firing (**Fig 12B**, **middle** trace) induced by the optogenetic stimulation which recovered (**Fig 12B, bottom** trace) after the washout of the drug. Again, the small-amplitude underlying depolarizations remained in the presence of the 4,9-anhydro-TTX in the 2 tested cases (**Fig 12B**, **middle,** top trace above the dotted baseline in inset).

## Discussion

The presented results provide evidence that astrocytes may regulate the excitability and firing patterns of primary afferent neurons in the trigeminal mesencephalic nucleus (NVmes). This regulation is mediated by a local astrocytic release of S100β along a specific axonal domain of NVmes neurons and the subsequent decrease of the extracellular calcium concentration ([Ca^2+^]_e_), causing *I*_NaP_ potentiation and repetitive firing.

### NVmes neurons properties and firing

The recorded neurons exhibited the distinctive electrophysiological properties previously described in the rat or mice NVmes (for review (33)): inward rectification upon membrane hyperpolarization (34, 35), accommodation of firing during depolarizing pulses (36), SMOs (10, 12), and burst generation dependent on persistent sodium current *I*_NaP_ (8, 9, 11, 12). These provide a recognizable signature for NVmes primary afferents, but in addition, we confirmed that all 160 neurons filled with Alexa 488 or 594 had the most typical unipolar morphology of primary afferents (37).

The vast majority (75%) of NVmes recorded neurons showed firing accommodation upon membrane depolarization. This intrinsic property of NVmes neurons relies on a 4AP-sensitive outward potassic current whose blockade causes repetitive firing (38). In our previous study in rat (39), the proportion of accommodating neurons was comparatively lower (64%). This difference between species has been reported by others (40) and is attributed to a significantly higher magnitude of the D-type K^+^ current in NVmes neurons from mice. Overall, as reported by others (25, 41), the NVmes neurons in our Na_V_1.6 knockout mice were less excitable than the ones from the WT mice since almost all (93%) of them showed firing accommodation. Emphasizing this decreased excitability, NVmes neurons from Na_V_1.6 null mice also exhibited a depolarised firing threshold relatively to WT NVmes neurons which was also reported for CA1 pyramidal cells (42).

Repetitive firing upon membrane depolarization and SMOs were observed in about a quarter of the total population of recorded NVmes neurons from the WT mice. However, more than half of 54 spike-adaptative neurons exhibited SMOs and repetitive firing with local applications of BAPTA or S100β, at some point along their axonal process. As already stated, SMOs and repetitive firing depend on *I*_NaP_ (8–10, 12) which is conveyed in these neurons by the Na_V_1.6 channel isoform (25). *I*_NaP_ is highly sensitive to the extracellular content of Ca^2+^ which reduces the current by occupying the pore of the channel (19). When Ca^2+^ is removed from the extracellular medium, closing of the Na^+^ channel slows or does not occur (19). The level of extracellular Ca^2+^ also modulates the voltage sensitivity of the gating kinetics of the Na^+^ channel (19). As calcium-chelating agents, it is not surprising that both BAPTA and S100β would increase the occurrence of SMOs and repetitive firing. By locally decreasing [Ca^2+^]_e_, they augment the probability of the Na_V_1.6 channels remaining in an open state, thereby increasing *I*_NaP_. Indeed, in a previous work conducted on the rhythmogenic neurons of the NVsnpr (20), both substances have been shown to increase the peak amplitude of the pharmacologically isolated *I*_NaP_ and to shift its activation and half-activation voltage towards more hyperpolarized potentials. The transition from adaptative to bursting-firing neurons with BAPTA and S100β is in agreement with Yang *et al.* (23) who showed that NVmes neurons excitability could be transformed from one class to another by manipulating the magnitude of *I*_4-AP_ and/or *I*_NaP_.

### [Ca^2+^]e chelation along NVmes neurons’ axonal processes elicits repetitive firing through NaV1.6 channel activation

Na_V_1.6 channels mostly localize in the distal end of the axon initial segment in neurons (17) and are presumed to be responsible for the action potential initiation (43, 44). Furthermore, using immunohistochemical labeling, Caldwell *et al.* (45) reported that Na_V_1.6 is also the predominant sodium channel at nodes of Ranvier in myelinated axons of the peripheral and central nervous systems. Our immunohistochemical data showed that the NVmes neurons possess Na_V_1.6 channels along their axons as well as in their cell bodies. This result is in agreement with the study of Chung *et al.* (46) which showed that the stem axons and somata of NVmes neurons are immunopositive to Na_V_1.6 antibody. This observed somatic immunoreactivity was weak, only located in the cytoplasm and could not be detected in the membrane. Given that *I*_NaP_ in these neurons mostly relies on the activation of Na_V_1.6 channels, the absence of these channels on the somata membrane would imply that NVmes neurons’ somata are devoid of *I*_NaP_. Indeed, inferring from the data acquired using pharmacological tools and dual patch recordings of the soma and the axon hillock of NVmes neurons, Kang *et al.* (47) concluded that NVmes neurons express different Na^+^ channels in the stem axon and the soma and, that *I*_NaP_ is likely present solely in the stem axon of these neurons. This could explain why there was no increase in excitability in the recorded neurons, in our study, when BAPTA and S100β were applied near their somata. However, 80% of the applications of BAPTA or S100β near the axonal process elicited firing in the mostly priorly quiescent NVmes cells. The elicited firing showed a significantly more hyperpolarised threshold potential than the firing evoked by standard step current injection that activates voltage-gated Na^+^ transient channels. This observation could be an argument in favor of these action potentials being generated via activation of Na_V_1.6 channels located on the axons of NVmes neurons since it is known that these channels display a hyperpolarized shift in their voltage-dependence of activation compared to other neuronal Na_V_s channels (48, 49) (reviewed in (50)). Accordingly, these firings were prevented with the selective blockade of the Na_V_1.6 channels with 4,9-TTX. Furthermore, in the Na_V_1.6 null mice, only 1 out of 22 applications (about 5 %) of both Ca^2+^-chelating agents near the axonal process of the NVmes neurons caused a firing response while the remaining applications mostly failed to trigger any kind of response which agrees with our immunohistochemical data revealing that, in the Na_V_1.6 knock-out mice, the axons of the NVmes neurons are immunonegative to Na_V_1.6. The immunopositivity to Na_V_1.6 that could be seen in the somata of the NVmes neurons in the Na_V_1.6 knock-out mice could be explained by the fact that the antibody used in our study targets a chain of amino acid residues located on the intracellular loop between domains II and III of the sodium channel α subunit (Alomone labs certificate of analysis for the anti-Na_V_1.6 (SCN8A) antibody; Cat #: ASC-009) which is not altered by the med mutation since it has been shown that the protein encoded by the med mutated gene is truncated within the first of the four transmembrane domains of the channel (29). Our immunohistochemical results in the Na_V_1.6 null mice suggest that the channel encoded by the mutated gene is not transported to the axon. This is on par with what has been reported by others (45).

In their study, Chung *et al*. (46) reported that mGluR-I expression was found to be colocalized with Na_V_1.6 on the stem axon of the NVmes neurons. They suggested that glutamatergic synaptic action on the stem axon would upregulate *I*_NaP_ at this location to trigger bursts. This could be a complementary mechanism as a source of firing. However, considering that no synaptic stimulation was applied in our study and that no spontaneous firing was observed in our recordings, it is more likely that the firings evoked were promoted by a direct effect of the punctual decreases of extracellular calcium on *I*_NaP_ along the axons of the NVmes neurons. Nevertheless, the proposed mechanism of a glutamatergic synaptic upregulation of *I*_NaP_ exerted on the axonal processes of NVmes neurons is quite interesting and may account for the increased SMOs and firings reported by Verdier *et al.* (39) in those neurons in response to electrical stimulation of surrounding trigeminal areas.

### A precise axonal subregion is involved in the Nav1.6-dependent firing

The AIS of NVmes neurons has not been overtly studied. Chung *et al.* (46) have images that suggest some enrichment of Na_V_1.6 channels in the stem axon emerging from the soma of NVmes neurons. However, the axons shown are not long enough to evaluate Na_V_1.6 channels further along the axon. Conversely, the AIS of the sensory neurons located in the DRG was studied by Nascimento *et al.* (51). They reported that in the TrkC-positive sensory neurons, which are the proprioceptive afferents located in the DRG and thus, represent the functional equivalent of most of the primary afferents whose somata are located in the NVmes, a 50-60 µm long initial segment, starting at 12 µm from the soma, could be observed in 25-30% of neurons. Considering that the Na_V_1.6 channels are absent in the more proximal part of the AIS (17), our firing responses with axonal applications of the Ca^2+^-chelating agents seem to agree with those morphological specifications since most of the firing responses that did not require facilitating depolarization were elicited with applications distanced 20-80µm from the soma. Although our physiological data seem to agree with the anatomical data reported for the AIS of the proprioceptive afferences located in the DRG, it is not clear whether the NVmes primary afferents possess an AIS since in Nascimento *et al.* (51) study, not all the TrkC-positive neurons showed an AIS.

### Anatomical relation between astrocytes and the NVmes neurons

In the periphery, satellite glial cells, which are the equivalent counterparts to the astrocytic glial cells of the central nervous system, closely envelop the cell bodies of the sensory neurons located within the DRGs and TGs, utterly isolating them from each other (reviewed in (52, 53). Rarely, does a cluster of sensory neurons share the same ensheathment (54). It was speculated that this peculiar organization allows for close interactions between SGCs and neurons (55) and emphasizes the absolute control these glial cells can exert over the extracellular milieu that surrounds these sensory neurons. Our immunohistochemical data reveal that although NVmes primary afferents are not as tightly enveloped by S100β-positive glial processes as DRG neurons, they are still well surrounded. S100β-positive glial cell bodies and processes are seen closely surrounding the somata and axons of these cells. However, the morphological relationship between astrocytes and NVmes neurons somata may be underestimated in our study because of our choice of astrocytic marker, since S100β mostly labels astrocytic cell bodies and proximal astrocytic processes. Using anti-GFAP antibodies in a study conducted on rats, Copray *et al.* (56) reported that each NVmes neuron was almost entirely sheathed with astrocytic processes radiating out from two or more astrocytes. Such anatomical organization places astrocytes in a strategic position to tightly regulate the excitability of the NVmes sensory neurons in the same way as the SGCs of the DRGs and TGs. Similarly, Serwanski *et al.* (57) reported that more than 95% of the nodes in the optic nerve, the corpus callosum, and the spinal cord of young adult mice or rats contained astrocyte processes. According to Dutta *et al.* (58), these close contacts with the Ranvier nodes along axons allow astrocytes to regulate the attachment of myelin to the axon through the vesicular release of thrombin protease inhibitors. They demonstrate that this regulation of the myelin attachment by astrocytes enables them to control the node structure, the myelin layer thickness, and ultimately, the speed of action potential conduction along the axon. However, independently of and in addition to this effect on myelin, this strategic arrangement of astrocytes at the nodes of Ranvier places them in a position where they can directly regulate the propagation of action potentials along the axon by releasing gliotransmitters that can act directly on the axonal membrane. The effect of released gliotransmitters could be all the more effective given the close apposition of glial processes and the neuronal membrane. Within the ganglia, the gap of extracellular space between the SGC sheath and the wrapped neuronal surface has a very constant distance, which is about 20 nm, comparable to the distance found in a synaptic cleft (reviewed in 53). The extracellular space at the synaptic cleft is so restricted that a change of just a few ions is enough to constitute a significant variation in concentration. This concept has been well modeled by Smith SJ. (60) who suggests that minimal variations in extracellular calcium at a synaptic cleft can correspond to major decreases in concentration and have a major effect on phenomena that depend on this concentration (59, 60). If a similar distance is found between the astrocytic endings and the axonal membrane of NVmes neurons, which is likely, given the close apposition revealed by our immunohistochemical data, the release of even a minute amount of S100β in such a confined space would produce a significant decrease of [Ca^2+^]_e_ around the nodes and powerfully activate the Na_V_1.6 channels located at the neurons’ nodes thus, facilitating the propagation of action potentials or even lead to the generation of ectopic discharges.

### Optogenetic activation of NVmes astrocytes along the axonal process of the primary afferent leads to ectopic firings through the release of S100β

Optogenetic activation of astrocytes via the expression of channelrhodopsin has been used by many and was shown to elicit intracellular calcium rises and to lead to the release of several gliotransmitters (21, 61–68). The specificity of the channelrhodopsin expression and the efficacity of the optogenetic stimulation on the illuminated astrocytes in the GFAP-ChR2 mice used in the present study have already been established in our more recently published work (21). Using immunofluorescence on brain tissue from the ChR2-EYFP mice, they found that the vast majority (94%) of V1 neuronal cell bodies were negative for EYFP, while in contrast, S100β-positive astrocytes expressed EYFP. They also assessed that the astrocytes were successfully activated by recording the membrane depolarizations and intracellular calcium increases evoked by the delivered optogenetic stimulation. Here, using this mouse line, we found that photostimulation of astrocytes surrounding the axon of NVmes neurons elicited firing that, as those evoked by local applications of BAPTA or S100β, relied on the activation of *I*_NaP_ and showed a significantly more hyperpolarised threshold potential than the firing evoked by current injection, suggesting that they may result from the release of S100β. Indeed, these firings could be reversibly blocked by application of an anti-S100β antibody. We did not test the effect of astrocytic photostimulation on [Ca^2+^]_e_, but according to Ryzcko *et al*. (21), such stimulation in the layer 5 of the visual cortex was associated with a drop of 0.13 mM. That a detectable drop could be recorded in a submerged preparation, continuously perfused with a calcium-loaded aCSF, is somewhat remarkable and we would presume that the calcium drop was likely greatly underestimated particularly in the extracellular space in area of close apposition between glial processes and neuronal membrane.

Photostimulation of NVmes astrocytes contributes to augmented neuronal excitability by amplifying SMOs and generation of ectopic repetitive firing that both rely on *I*_NaP_. The unique pseudounipolar morphology of these centrally located neurons points to this regulatory effect of astrocytes taking place along the axonal process and adds to the accumulating evidence that these glial cells interfere with signal processing at all levels of the input-output computation within neuronal circuits. Involvement of *I*_NaP_ in distal axon excitability has already been demonstrated in CA1 pyramidal neurons (69) but the present study supplements this demonstration by pointing astrocytes as the culprit of *I*_NaP_ regulation along axonal process through the release of S100β.

### Functional considerations

#### Synchronization and signal amplification

NVmes neurons are known to be electrically coupled by gap junctions (40, 70–72). But, while astrocytes are reported to form extended networks, in diverse brain areas (73–76) that are often function-related (75, 76), coupling within the NVmes is mostly restricted to pairs or small clusters of neurons linked by Cx36 at cell-cell appositions (71). However, despite the small number of coupled neurons, Curti *et al.* (71) found that electrical coupling between NVmes neurons is strong and promotes robust spike synchrony among pairs of these afferents through a synergistic interaction with those neurons’ active membrane properties. The authors also show, using a combination of Neurobiotin injection and Cx36 labeling, that in many cases a lack of tracer-coupling was observed between adjacent neurons displaying Cx36 labeling at appositions suggesting that the channels formed by Cx36 between these neurons could be closed. In fact, they estimated that coupling between NVmes neurons is supported by a very small proportion of the gap junction channels and suggested that it could be under regulatory control. Indeed, coupling between cells is dynamic and can be up or downregulated. The [Ca^2+^]_e_ is one of the factors that can greatly influence gap-junction coupling (77). Therefore, even if the cell body of NVmes neurons lacks *I*_NaP_, the S100β released by the surrounding astrocytes may contribute to an increased excitability and synchronized firing by allowing the transfer of the antidromic action potentials generated along the axon to a large number of cells through upregulation of gap-junction coupling by modulating extracellular calcium. This astrocytic control of coupling is of particular interest in a nucleus where the cells are majorly refractory to discharge since it may allow to bypass these neurons refractory nature by coupling the more excitable cells to the refractory ones. In our study, 43% of the recorded neurons fire a single action potential in response to 1 s long depolarizing pulses current injection. According to Dapino *et al*. (40), mice NVmes neurons are more refractory to firing than rat NVmes neurons due to a higher expression of a 4-AP-sensitive outward K^+^ current (D-type) that counteracts postsynaptic recruitment through electrical synapses. However, action potentials ectopically generated in the axon are likely to invade the soma and release the strong inhibitory control exerted by this conductance. Interestingly, in this respect, Saito *et al.* (78) reported that 4-AP-highly sensitive K^+^ channels are located exclusively on the somata of NVmes neurons. These channels, according to their electrophysiological data, suppress spike initiation by increasing firing threshold, and then may facilitate successive spike invasion by shortening the refractory period, thus favoring distally initiated action potentials. The ensuing depolarization would then presumably inactivate I_4-AP_ and allow action potential transfer from cell to cell through the electrical coupling.

Davoine and Curti (79) proposed that the reciprocal interaction that exists between active electrical properties and electrical coupling endows circuits of coupled neurons with the capability to act as coincidence detectors and since astrocytes act on both parameters, we propose that they may play an important role in this neuronal function as well.

Lastly, independently of the coupling between NVmes neurons, it is known that a single astrocyte, because of its highly ramified morphology, contacts several cells simultaneously. Thus, a concerted action of a single astrocyte on the nodes of adjacent axons may suffice to elicit a synchronized activity between many neighbouring cells. Pondering about the implications of Sasaki *et al.* (80) study, showing that stimulation of peri-axonal astrocytes causes broadening of action potentials in axons of CA3 pyramidal neurons, Debanne and Rama (81) postulated that since a single astrocyte covers a spherical volume with a diameter of ∼40 μm, the activity of an astrocyte may affect a large population of axons, creating volumes of enhanced synaptic transmission. If we add to this that several astrocytes can be coupled together to form a functional unit as in the NVsnpr (82), that brings the influence of astrocytes to another level and gives a glimpse of the powerful control they may exert on the coordination of firing across networks of neurons.

#### Neuronal compartmentalization

In their study about the involvement of *I*_NaP_ in distal axon excitability of CA1 pyramidal neurons, Muller *et al*. (69) reported that the contribution of *I*_NaP_ to excitability was not equal in all axonal branches, implying that all branches are not subjected to the same control. Distinctive inputs to separate parts of a neuron sometimes contribute to neuronal compartmentalization which has been suggested to occur to NVmes neurons during fictive mastication (83). Westberg and collaborators (83) reported that during the jaw-closing phase, antidromic action potentials appear to be generated in the central axons of the NVmes neurons. These action potentials were not detected in the soma and stem axon area and their generation was imputed to strong phasic GABA-mediated PADs since they were found in axonal branches located in areas containing populations of interneurons that fire rhythmically during fictive mastication. The authors proposed that during jaw closure, the central axon becomes functionally disconnected from the stem axon and soma, which continue to transmit signals from muscle spindles with little antidromic contamination, to act as a premotoneuron carrying inputs from the masticatory CPG to motoneurons. Although the generation of action potentials by PADs is a likely mechanism, it’s also possible that the antidromic potentials generated could result from the release of S100β by astrocytes surrounding the central axon branches in these regions. The NVmes neurons’ central axon is quite extensive and in jaw muscle spindle afferents, the central axons’ collaterals reach several brainstem areas (37, 84, 85) that do not necessarily need to receive the same undisturbed signal from the periphery. It is therefore highly probable that the masticatory CPG regulates the information relayed by the various collaterals by inhibiting or amplifying it, and astrocytes could be an effective player in this control.

#### Pathologies

There is accumulating evidence that the SGCs in DRG are activated in chronic pain conditions (reviewed in (53)). It has been proposed that these activated SGCs contribute to the establishment and maintenance of chronic pain. However, a specific mechanism for this contribution has yet to be determined. The astrocytes of the NVmes also become reactive in the acidic saline masseteric myalgia model (86) in which NVmes neurons showed increased excitability (22). Reactive astrocytes are known to release increased amounts of gliotransmitters (87). Particularly, regarding S100β, it has been shown that S100β mRNA and protein are increased in the spinal cord after peripheral inflammation and nerve injury (88). Also, increased levels of serum S100β have been reported in diverse chronic pain conditions (89–91). It is therefore possible that reactive astrocytes contribute to this increased excitability of the NVmes neurons in this pain model by releasing excessive amounts of S100β in the vicinity of nodes and demyelinated portions of the axon. This mechanism may be at play in any pathology where neuronal hyperexcitability or hypersynchrony is reported.

### Concluding remarks

Others have reported regulation of axonal excitability by astrocytes directly, through purinergic (92) and glutamatergic (80) signaling, or indirectly, by acting on the myelin (58). It therefore appears that astrocytes have a variety of mechanisms at their disposal to impact the electrical properties of axons. In this study, we showed how astrocytes, by releasing S100β in the vicinity of a specific segment of the axonal process of NVmes neurons modulate their excitability by acting on *I*_NaP_. Those very peculiar sensory neurons have classically been associated with chewing, an essential rhythmic behavior sustained by an extensive brain circuit (reviewed in (93)). However, there is accumulating evidence that their functions go beyond the relaying of jaw proprioceptive inputs for oro-sensory–motor control. Indeed, those cells are involved in numerous other brain functions, such as whisker pad proprioception (94, 95), food intake regulation (96), stress-induced masseter hyperactivity (97) and sleep physiology (98), to name a few. This implies that understanding the factors that regulate their excitability is paramount, and more studies are required to reach this objective. The present study clearly shows that unraveling the involvement of astrocytes in this control is a promising avenue.

## Material and methods

All experiments were conducted according to the Canadian Institutes of Health Research rules and were approved by the Animal Care and Use Committee of Université de Montréal.

A total of 117 mice were used, including 44 wild-type mice, (C57BL/6, Charles River, Wilmington, Massachusetts, USA), 5 Nav1.6 null mice, and 68 mice expressing the channelrhodopsin 2 (ChR2) under the control of the GFAP promoter (GFAP-ChR2-EYFP mice). Nav1.6 null mice were obtained by crossing heterozygous Scn8a^med^ mice (C3Fe.Cg-Scn8a^med^/J, stock 003798, JAX) and selecting the homozygous offsprings. GFAP-ChR2-EYFP mice were produced by crossing GFAP-Cre (B6.Cg-Tg(Gfap-cre)73.12Mvs/J, stock 12886, JAX) and ChR2-lox mice (B6.Cg-Gt(ROSA)26Sortm32(CAG COP4∗H134R/EYFP)Hze/J, stock 24109, JAX).

### Brainstem slice preparation

Coronal brainstem slices (325-350 µm) were prepared from mice aged from 14 to 21 days. The mice were anesthetized by inhalation of isoflurane (Pharmaceutical Partners of Canada Inc., Richmond Hill, ON, Canada) prior to decapitation. Their brain was quickly extracted from the cranium and sectioned in an ice-cold modified artificial cerebrospinal fluid (CSF, in mM: 3 KCl, 1.25 KH2PO_4_, 4 MgSO_4_, 26 NaHCO_3_, 10 Dextrose, 0.2 CaCl_2_, 219 Sucrose, pH 7.3-7.4, 300-320 mOsmol/kg) saturated with a mix of 95% O_2_ and 5% CO_2_ using a VT1000S vibratome (Leica). Slices were transferred to a submerged chamber and continuously perfused with artificial CSF (in mM: 124 NaCl, 3 KCl, 1.25 KH2PO_4_, 1.3 MgSO_4_, 26 NaHCO_3_, 10 Dextrose, and 1.6 CaCl_2_, pH 7.3-7.4, 294-300 mOsmol/kg) bubbled with 95% O_2_ and 5% CO_2_ at room temperature. Slices were allowed to rest for a minimum of one hour before recording.

### Electrophysiology and analysis

Recordings were carried out at room temperature in a submerged chamber continually perfused with artificial CSF bubbled with 95% O_2_ and 5% CO_2_. Patch microelectrodes were pulled from borosilicate glass capillaries (1.5 mm outside diameter, 1.12 mm inside diameter, World Precision Instruments) using a P-97 puller (Sutter Instruments). For neuronal recordings, pipettes (resistance 6-10 MΩ) were filled with an internal solution containing (in mM): 140 K-gluconate, 5 NaCl, 2 MgCl_2_, 10 HEPES, 0.5 EGTA, 2 Tris ATP salt, 0.4 Tris GTP salt, pH 7.2-7.3, 280-300mOsmol/kg. 0.05 Alexa Fluor 488 or 594 was added to the internal solution to visualize neuronal and axonal morphologies during the experiment. Confocal imaging was performed using an Olympus Fluoview FV 1000 confocal microscope equipped with a 40x (N.A. 0.80) water immersion objective. All recordings were performed using a Multiclamp 700A amplifier, Digidata 1322A interface coupled to a computer equipped with pClamp 8 software (Molecular Devices, San Jose, CA). After the establishment of a gigaseal, the membrane potential was held at -60 mV, and the membrane patch was suctioned. The pipette resistance and capacitance were compensated electronically. Neurons were discarded when action potentials did not overshoot 0 mV or when the resting membrane potential was depolarized (>-45 mV). Recordings were analyzed using standard scripts in Clampfit. The neuronal passive properties included the resting membrane potential (in mV), the input resistance (in MΩ), and the firing threshold. The resting membrane potential was measured when no current was injected into the recorded neuron in current-clamp mode. The input resistance was determined as the voltage change induced by a small hyperpolarizing current (−20 to−40 pA) of 1 s duration applied from resting membrane potential divided by the amount of injected current. Firing patterns and membrane rectification were described from the current-voltage (I-V) curve obtained in the current-clamp mode.

### Optogenetic stimulation

Astrocytes were optogenetically stimulated with two lasers (440 and 488 nm) used simultaneously in the SIM lightpath of an FV1000 microscope (Olympus) in mice expressing the channelrhodopsin (ChR2) under the control of the GFAP promoter (GFAP-Cre/ChR2-lox). The SIM scanner was used in the ‘tornado’ scanning mode (a spiraling scan mode) to photoactivate manually delineated small areas surrounding the recorded neuron. Optogenetic stimulations were applied using 5-30 s pulses (10-20% laser power/8.6-14.9 µW for laser 440 nm/8.7-15.9 µW for laser 488 nm). To assess the specificity of wavelength in activating astrocytes, 559 nm laser pulses (10-20% laser power/120-200 µW) were applied in control experiments.

### Drug application

Chemicals used in this study were purchased from Sigma-Aldrich (Oakville, Ontario, Canada), Tocris Biosciences (Ellisville, Missouri, USA), Abcam (Cambridge, UK), and Inixium (Laval, Quebec, Canada). The following drugs were bath-applied using a syringe pump: 4,9-anhydro-tetrodotoxin (4,9-anhydro-TTX, 100 nM), CNQX (10 µM), D,L-2amino-5 phosphonovaleric acid (APV, 75 µM), SR 95531 hydrobromide (Gabazine, 20 µM). In some experiments, the Ca^2+^-binding proteins S100β (129 μM) or 1,2-bis(o-aminophenoxy)ethane-N,N,N′,N′-tetraacetic acid tetrasodium salt (BAPTA, 5 mM) were locally applied with glass micropipettes (tip diameter around 1 μm) with 2-20 psi pressure pulses of variable duration (1-30 s; Picospritzer III, Parker Instrumentation, Fairfield NJ USA). Monoclonal anti-S100β antibodies (mouse anti-S100β, Sigma Aldrich #S2532 or rabbit anti-S100β, Abcam ab56642) were applied locally with large-tip (10-20 µm) glass micropipettes carefully lowered near the recorded neuron with 0.1-2 psi pressure pulses lasting from 5 to 20 minutes.

### Immunohistochemistry

Coronal brainstem slices (500 µm) were prepared from 14 to 21-day-old wild-type, Na_v_1.6-null, or GFAP-cre mice using a vibratome VT 1000S (Leica), immediately immersed in a solution of 4% (wt/vol) paraformaldehyde in PBS and kept overnight at 4°C. For cryoprotection, the slices were then immersed in a solution of 20% sucrose in PBS for two hours at 4°C. Using a sliding microtome (Leica SM20000R), 40 µm thick sections of the brainstem were made. The sections were rinsed 3 times for 10 minutes in PBS and incubated in a blocking solution containing 0.3% Triton X-100 and 10% Normal Donkey Serum (Jackson ImmunoResearch #017-000-121) in PBS for two hours at room temperature. The sections were then rinsed 3 times for 10 minutes in PBS and incubated overnight at 4°C in a mix of the primary antibodies (mouse anti-S100β, Sigma Aldrich #S2532, dilution 1:400; rabbit anti-Nav1.6, Alomone lab #ASC-009, dilution 1:400; chicken anti-MBP, ThermoFisher PA1-10008, 1:500; or sheep anti-Parvalbumin, ThermoFisher, PA47693, dilution 1:40) diluted in the blocking solution. The following day, the slices were rinsed 3 times for 5 minutes in PBS and incubated in the relevant secondary antibodies mix (donkey anti-sheep Alexa Fluor 405, Jackson Immunoresearch #713-475-003, dilution 1:500; donkey anti-rabbit-Alexa Fluor 488, Jackson Immunoresearch #711-545-152, dilution 1:500; donkey anti-mouse Alexa Fluor 488, Jackson Immunoresearch #715-545-151, dilution 1:500; donkey anti-rabbit Alexa Fluor 555, Invitrogen #A31572, dilution 1:500; donkey anti-mouse-Alexa Fluor 594, Jackson Immunoresearch #715-585-150, dilution 1:500; or donkey anti-chicken Alexa Fluor 647, Jackson Immunoresearch #703-605-155, dilution 1:500) diluted in the blocking solution for 60 minutes in a dark chamber at room temperature. The sections were then rinsed 3 times for 5 minutes in PBS and mounted on ColorFrost Plus slides (Fisher Scientific, Ottawa, Ontario, Canada) using Fluoromount-G (Southern Biotech, Birmingham, Alabama, USA). Slides imaging was done using either an E600 epifluorescence microscope equipped with aDXM1200 digital camera (Nikon), an FV1000 confocal microscope (Olympus), or a TCS SP8 STED nanoscope (Leica). In all cases, a negative control was performed by removing the primary antibody and the absence of specific labeling on brainstem sections was confirmed. Images were treated with ImageJ (NIH) software to combine pictures and adjust the levels so that all fluorophores were clearly visible simultaneously.

### Statistical analysis

Data are presented as mean ± standard error to the mean (SEM) and as proportions (%). Our sample sizes are comparable to those employed in the field and were not predetermined by any statistical methods. Our experimental paradigms necessitated no blinding or randomization procedures. For pairwise comparisons of normally distributed data, paired *t*-tests or independent *t*-tests were used. For pairwise comparisons of non-normally distributed data, the Wilcoxon Signed-rank, or the Kruskall Wallis tests were used. Comparisons of percentages were done with χ^2^ tests. Statistical significance was defined as *P* < 0.05. Data analysis was performed using the statistical Sigma Stat software and SPSS.

## Acknowledgments

Rafael Sanz Galvez generously performed the immunostaining of S100β and Pvalb in the NVmes.

